# Intrinsic DNA codes govern distinct modes of nucleosome-transcription factor interactions

**DOI:** 10.64898/2025.12.30.697010

**Authors:** Christopher W. Carson, Sheethal Umesh Nagalakshmi, Image Adhikari, Julia M. Freewoman, Joseph R. Pizzi, Prakyat Prakash, Feng Cui

## Abstract

The interplay between transcription factors (TFs) and nucleosomes is central to gene regulation, yet most studies adopt a protein-centric perspective that largely overlooks DNA sequence contributions. Here, we identify four distinct nucleosomal DNA classes, including the canonical WW/SS pattern (W = A/T, S = G/C). All four nucleosome types are widespread across the human genome both *in vivo* and *in vitro*. Nucleosomes sharing the same pattern are rotationally in phase, whereas different patterns exhibit specific rotational offsets, revealing distinct DNA-encoded codes for nucleosome positioning. Analysis of 747 TFs across eukaryotes shows that the WW/SS and anti-WW/SS patterns are strongly associated with TF binding in chromatin. Differences in the proportions of these two patterns, quantified as ΔNPS values, correlate closely with *in vitro* nucleosome occupancy. Integrating ΔNPS with *in vivo* nucleosome occupancy uncovers four distinct modes of nucleosome-TF interaction and provides new insight into nucleosome-depleted regions around TF binding sites.

## Introduction

Understanding the interaction modes between transcription factors (TFs) and nucleosomes is central to gene regulation^1–3^. The traditional view holds that TFs compete with nucleosomes for access to their binding sites^4,5^, a model supported by ChIP-seq analyses showing that TF-bound genomic regions are typically nucleosome-depleted regions (NDRs)^6^. However, numerous studies have demonstrated that a subset of TFs, including pioneer factors, can directly bind nucleosomal DNA^7,8^.

Most studies of nucleosome-TF interactions have relied on protein-centric experimental approaches, such as electrophoretic mobility shift assays^9,10^, single-molecule fluorescence methods^11,12^, cryogenic electron microscopy^13–15^, SELEX^16^, and protein microarray^17^. These experiments typically employed a limited number of nucleosomal DNA templates, including the synthetic ‘601’ sequence^18^ or a small set of genomic loci with strong nucleosome positioning sequences (NPSs), such as yeast Hsp82^19^, ribosomal protein L30 genes^20^, mouse ALBN1, NRCAM, and CX3CR1^17^, and the human lin28b locus^21^. Consequently, prior work has focused primarily on TFs rather than their DNA binding partners.

Nucleosome-TF interactions are influenced by the rotational phasing of TF binding motifs within nucleosomes^22–25^. Early studies of the glucocorticoid receptors showed that motif accessibility depends on rotational positioning^22^, with similar effects observed for p53^23,24^ and SRY-box 2^25^. Rotational positioning is defined by which face of the DNA duplex contacts the histone core^26^ and is strongly determined by DNA sequence^27–33^.

One of the best-characterized sequence motifs is the WW/SS pattern (W = A/T, S = G/C), in which WW dinucleotides preferentially occupy minor-groove bending sites (minor-GBS) and SS dinucleotides occupy major-groove bending sites (major-GBS)^34^. This pattern exhibits a ∼10-bp periodicity and is structurally stabilized by conserved “sprocket” arginine residues that insert into minor grooves^35–37^, favoring interactions with AT-rich DNA that narrows the minor groove^38–40^. These periodic interactions form the structural basis for rotational positioning.

An opposite sequence motif, termed anti-WW/SS, places WW and SS dinucleotides in reversed groove contexts^41^, leading to less favorable histone–DNA interactions. We previously showed that anti-WW/SS nucleosomes are enriched in mammalian genic regions^42^ and potentially stabilized by H4 and H2A histone N-tails^43^, but their role in nucleosome-TF interactions remained unclear.

Here, we identified four distinct nucleosomal DNA patterns in the human genome, Types 1-4. Nucleosomes sharing the same pattern are rotationally in phase, whereas different types exhibit specific phasing relationships. We show that Type-1 and Type-4 nucleosomes are preferentially associated with TF binding. The difference in their regional abundance, quantified as ΔNPS, strongly correlates with *in vitro* nucleosome occupancy (NO). Integrating ΔNPS with *in vivo* NO reveals multiple modes of nucleosome-TF interaction.

## Results

### Four types of nucleosomes are widespread across the human genome

We extracted nucleosome core particle (NCP) DNA from a human *in vitro* nucleosome map^44^ and an *in vivo* map generated in GM12878 cells^45^ (Supplementary Table S1). As shown previously^42^, NCP fragments of 147 bp can be classified into four sequence patterns, Types 1-4, based on the relative frequencies of WW and SS dinucleotides across the 12 minor- and 12 major-GBS (Figure 1A). Type-1 nucleosomes exhibit the canonical WW/SS pattern, with WW enriched in minor-GBS and SS enriched in major-GBS. Type-2 and Type-3 nucleosomes display “mixed” patterns: Type-2 nucleosomes show higher WW and SS occurrence in minor-GBS than in major-GBS, whereas Type-3 nucleosomes show the opposite. Type-4 nucleosomes represent the inverse of the WW/SS pattern, with WW enriched in major-GBS and SS enriched in minor-GBS (Figure 1B) and are therefore referred to as anti-WW/SS nucleosomes.

**Figure 1.**
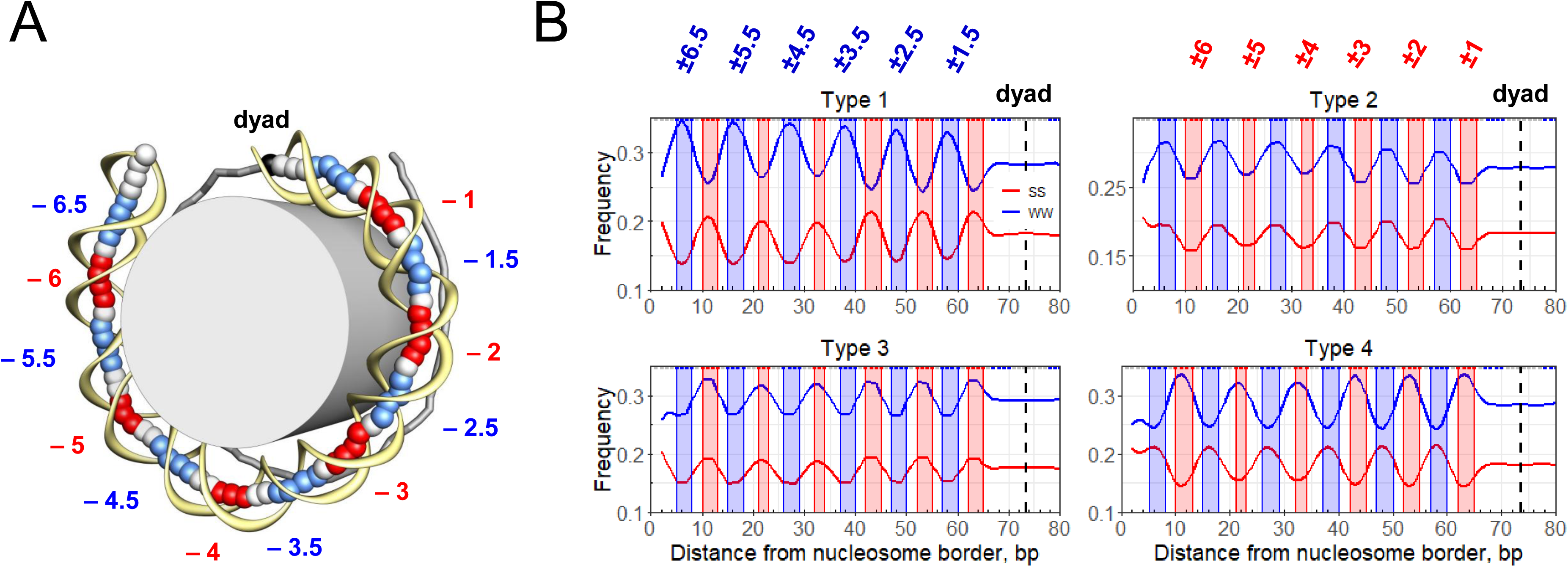
Minor- and major-groove bending sites and four WW/SS-based sequence patterns along nucleosomal DNA. (A) Schematic representation of the nucleosome core particle (PDB: 1KX5) containing 147 bp of DNA. The DNA duplex is divided into two halves separated by the dyad (diamond). In the “ventral” half of the particle, base-pair centers are shown as large spheres, and the sugar–phosphate backbone is rendered as a yellow ribbon. In the “dorsal” half, sticks connect base-pair centers (only small segments are visible). minor-GBS and major-GBS are highlighted in blue and red, respectively, and are labeled according to their superhelical locations (SHLs). (B) Four nucleosomal DNA sequence patterns identified in GM12878 cells (*in vivo*). At each nucleosomal position, the frequencies of AA, TT, AT, and TA dinucleotides (WW, shown in blue) and GG, CC, GC, and CG dinucleotides (SS, shown in red) were computed and symmetrized relative to the dyad (dashed line). Three–base pair running averages for WW and SS frequencies are plotted for Type-1, Type-2, Type-3, and Type-4 nucleosomes. Blue and red shaded regions denote the minor- and major-GBS, respectively, corresponding to their SHL designations. Type-1 nucleosomes correspond to the canonical WW/SS pattern, whereas Type-4 nucleosomes correspond to the inverted anti-WW/SS pattern.

All four nucleosome types are present in both *in vivo* (Figure 1B) and *in vitro* (Supplementary Figure 1) maps. In both datasets, Type-1 nucleosomes are the most abundant class, followed by Type-4, Type-2, and Type-3 nucleosomes (Supplementary Tables S2–S3). Comparing their relative fractions reveals notable differences between *in vitro* and *in vivo* landscapes. The *in vitro* map contains ∼3% more Type-1 nucleosomes and ∼3% fewer Type-4 nucleosomes compared to the *in vivo* map, whereas the differences for Type-2 and Type-3 nucleosomes are much smaller (∼1%). This suggests that Type-1 nucleosomes are energetically favored *in vitro*, likely because nucleosome positioning in the absence of chromatin-associated proteins reflects intrinsic histone-DNA preferences.

These observations imply that chromatin-associated proteins, such as transcription factors, chromatin remodelers, and RNA polymerase, redistribute nucleosomes with different DNA-encoded patterns. Notably, this redistribution exerts stronger effects on Type-1 and Type-4 nucleosomes than on Type-2 and Type-3. Consistently, the difference in abundances between Type-1 and Type-4 nucleosomes is 9.8% *in vitro*, nearly three-fold higher than the corresponding difference *in vivo* (3.3%). In contrast, the shift between Type-2 and Type-3 nucleosomes is more modest (7.6% *in vitro* versus 5.6% *in vivo*, Supplementary Table S4).

In summary, two key conclusions emerge. First, all four nucleosome types are broadly distributed across the human genome in both *in vitro* and *in vivo* conditions. Second, Type-1 and Type-4 nucleosomes exhibit substantially larger differences between *in vitro* and *in vivo* contexts than Type-2 and Type-3, indicating that chromatin proteins exert stronger regulatory influence on nucleosomes with these two sequence patterns.

### The four types of nucleosomes exhibit distinctive genomic organization

To investigate how the four nucleosome types are organized across the genome *in vivo* and *in vitro*, we first computed normalized nucleosome occupancy at every base pair and then calculated pairwise correlations between nucleosome maps. Both *in vivo* and *in vitro* datasets revealed a consistent pattern. The full nucleosome map showed the highest correlation with the Type-1 map (0.74 *in vivo*, 0.76 *in vitro*; Figure 2A and Supplementary Figure S3A), consistent with the fact that Type-1 nucleosomes are the most abundant class (Supplementary Table S4). The second-highest correlation was observed between the full map and the Type-4 map (0.72 *in vivo*, 0.71 *in vitro*; Figure 2D and Supplementary Figure S3D). Lower correlations were observed for Type-2 (0.60 *in vivo*, 0.63 *in vitro*; Figure 2B) and Type-3 (0.45 *in vivo*, 0.54 *in vitro*; Figure 2C), reflecting their lower abundance. Notably, the ranking of these correlations mirrors the relative frequencies of the four nucleosome types (Supplementary Table S4).

**Figure 2.**
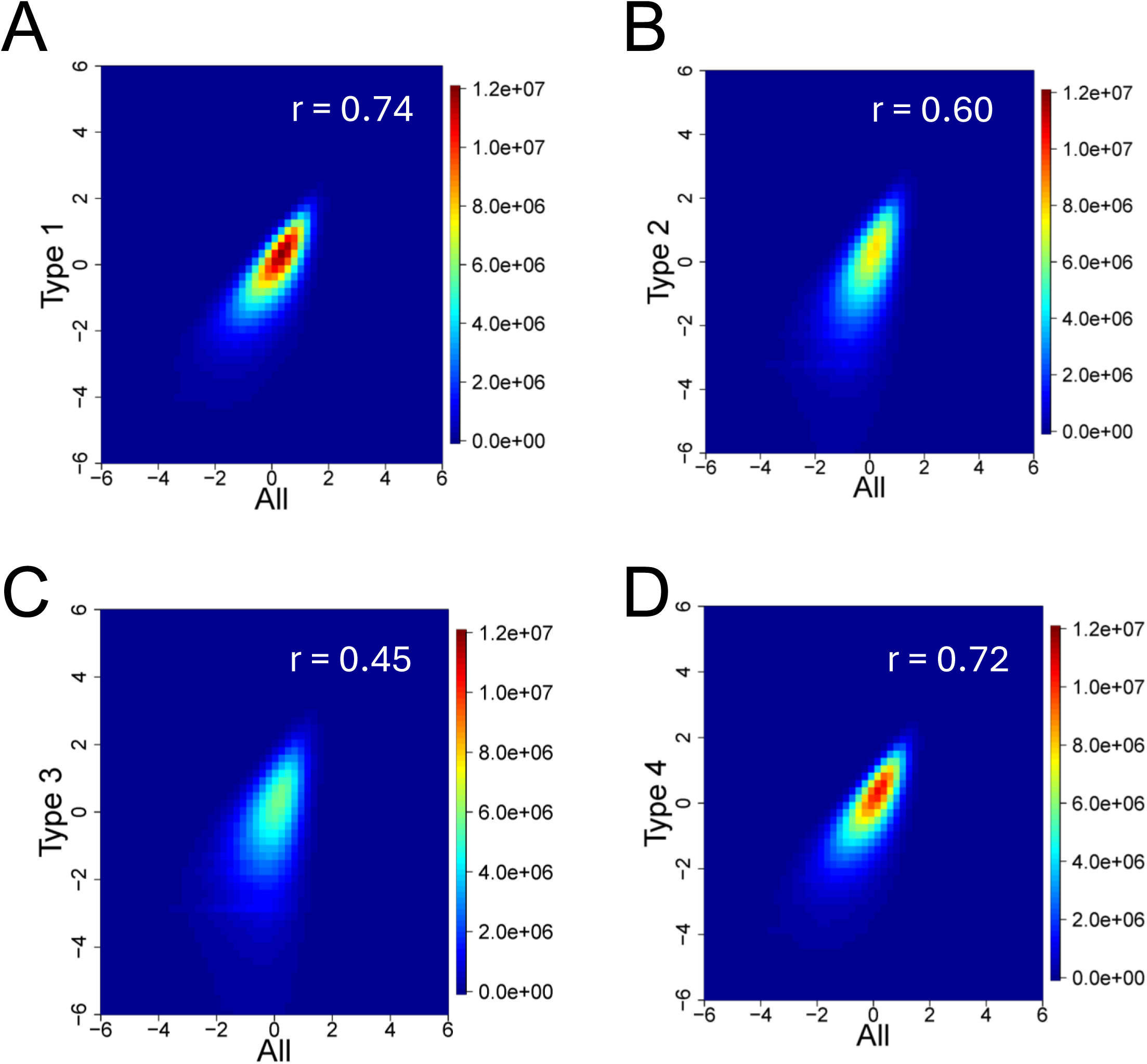
Comparison of *in vivo* nucleosome occupancy maps across the four nucleosome types. Density dot plots show normalized nucleosome occupancy per base pair in the complete nucleosome dataset plotted against occupancy in each nucleosome type: Type-1 (A), Type-2 (B), Type-3 (C), and Type-4 (D). The color scale reflects the number of data points within each region of the plot. Pearson correlation coefficients were computed for each pairwise comparison to quantify the similarity between the full nucleosome map and individual nucleosome-type maps.

Comparisons among the four nucleosome-type maps yielded several additional insights. First, Type-1 and Type-4 nucleosome maps exhibited the strongest cross-correlation (0.51 *in vivo*, 0.47 *in vitro*; Supplementary Figures S2C and S3G). Second, Type-2 and Type-3 maps showed the next highest correlation values (0.41 *in vivo*, 0.39 *in vitro*; Supplementary Figures S2D and S3H). Third, all remaining pairwise comparisons displayed relatively low correlations, ranging from 0.13–0.24 *in vivo* (Supplementary Figures S2A–B, S2E–F) and 0.27–0.32 *in vitro* (Supplementary Figures S3E–F, S3I–J).

Together, these results indicate that nucleosomes with certain DNA sequence patterns tend to co-localize across the genome. Specifically, Type-1 and Type-4 nucleosomes appear to occur together, whereas Type-2 and Type-3 nucleosomes form a separate organizational group. This suggests that the four nucleosome types exhibit distinct spatial distributions along chromatin, likely reflecting the DNA-encoded rotational preferences and their interactions with chromatin-associated factors.

### Nucleosomes with different sequence patterns have specific rotational phasing

To examine the relative positioning of the four nucleosome types across the genome, we applied the distance auto-correlation (DAC) and distance cross-correlation (DCC) functions described previously^46^. The DAC function measures the positional relationships between pairs of NCPs from the same set (e.g., all Type-1 nucleosomes), whereas the DCC function measures positional relationships between NCPs from two different sets (e.g., Type-1 versus Type-4 nucleosomes).

We first computed the DAC function for the complete nucleosome set, which contains all four sequence patterns. In the *in vitro* dataset, weak peaks appear approximately at 10, 20, 30, and 40 bp (Supplementary Figure S4), consistent with earlier reports^44^. These ∼10-bp periodicities reflect overlapping nucleosome positions that share the same rotational orientation, indicating that DNA is wrapped such that the helical face contacting the histone octamer repeats every full turn of the DNA helix. In contrast, the *in vivo* dataset showed no such periodicity (Supplementary Figure S4), suggesting that the intrinsic rotational pattern can be disrupted by chromatin-associated proteins.

We then computed the DAC function for each nucleosome type separately. Strikingly, all four nucleosome types exhibited strong ∼10-bp periodicity *both in vivo and in vitro* (Figure 3A–D). Type-1 nucleosomes showed clear peaks at ∼10-bp intervals (Figure 3A), as did Type-2 (Figure 3B), Type-3 (Figure 3C), and Type-4 (Figure 3D) nucleosomes. These patterns indicate that nucleosomes sharing the same sequence pattern are rotationally in phase with one another, regardless of cellular context.

**Figure 3.**
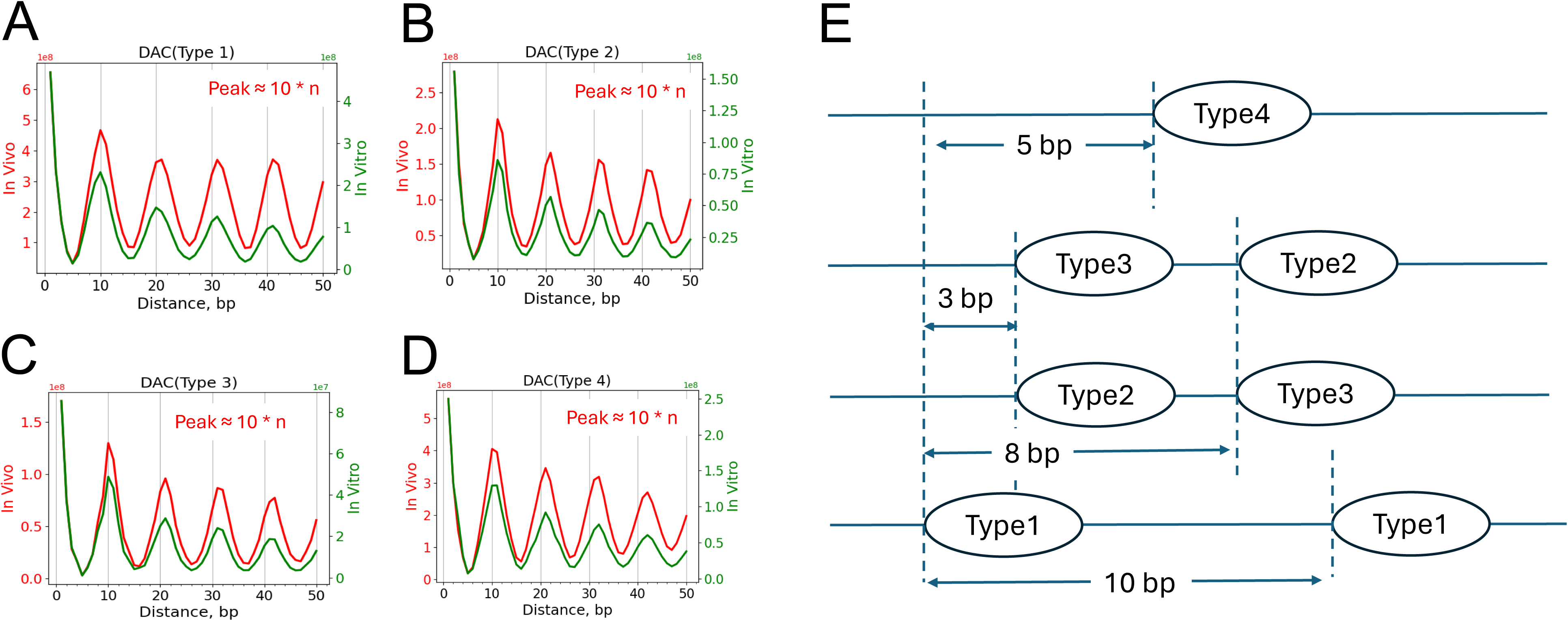
Distance auto-correlation profiles and a model for the relative positioning of the four nucleosome types. (A–D) Distance auto-correlation (DAC) functions for Type-1 (A), Type-2 (B), Type-3 (C), and Type-4 (D) nucleosomes in *in vivo* (red) and *in vitro* (green) datasets. Peaks occur approximately at distances of 10 × *n* bp (where *n* = 1, 2, 3, …), indicating ∼10-bp periodicity and in-phase rotational alignment among nucleosomes of the same type. (E) Model illustrating the relative positioning of the four nucleosome types. Dashed lines represent the start positions of nucleosomes. The predicted spacing relationships among nucleosome types are derived from the peak locations of the DAC functions (A–D) and from DCC analyses (Supplementary Figure S5).

Analysis of the DCC function between different nucleosome types yielded several additional insights. First, DCC peaks were generally stronger in the *in vivo* dataset than *in vitro* (Supplementary Figure S4), likely to reflect the larger number of the *in vivo* NCPs (Supplementary Tables S2-S3). Second, Type-1 and Type-4 nucleosomes displayed strong out-of-phase peaks at ∼5, ∼15, ∼25, ∼35, and ∼45 bp (Supplementary Figure S5C) in both datasets, indicating a consistent half-turn rotational offset. Third, the same out-of-phase relationship was observed between Type-2 and Type-3 nucleosomes (Supplementary Figure S5D). Finally, all other pairwise comparisons produced two sets of peaks, one at ∼13, ∼23, ∼33 bp and another at ∼18, ∼28, ∼38 bp, across multiple type pairs (Supplementary Figure S5A–B, E–F), further demonstrating that nucleosomes with distinct sequence patterns have characteristic rotational spacing.

Because the sum of all individual DAC and DCC functions recapitulates the DAC function of the complete nucleosome population (Supplementary Table S5-S6), these patterns reveal that nucleosomes with different sequence motifs are arranged in a highly regular but previously hidden rotational organization.

Based on these results, we propose a rotational phasing model for the four nucleosome types, using Type-1 nucleosomes as the reference (Figure 3E). Type-1 nucleosomes are positioned ∼10 bp apart due to their in-phase relationships. Type-4 nucleosomes, which are out of phase with Type-1, are offset by ∼5 bp. For Type-2 and Type-3 nucleosomes, Type-1 nucleosomes are separated by distances of either 10n + 3 bp or 10n + 8 bp (where *n* = 0, 1, 2, …), consistent with the two peak groups observed in the DCC analyses (Supplementary Figure S5A–B). Type-2 and Type-3 nucleosomes themselves are ∼5 bp apart, reflecting their out-of-phase relationship (Supplementary Figure S5D).

In summary, we identify four distinct nucleosomal DNA sequence patterns that serve as DNA-encoded rotational positioning codes across the human genome. Nucleosomes with the same sequence pattern align rotationally in phase, whereas nucleosomes with different patterns adopt specific, predictable rotational offsets relative to one another.

### Type-1 and Type-4 nucleosomes are associated with TF binding

Because Type-1 and Type-4 nucleosomes show the largest differences between *in vitro* and *in vivo* maps (Supplementary Table S4), we hypothesized that these two nucleosome types influence TF binding *in vivo*, and that such effects should be reduced at motifs not bound by TFs. We used CTCF as a model system because its nucleosome organization has been extensively characterized^47^.

We began by examining nucleosome occupancy around CTCF motifs. As expected, *in vivo* NO shows a characteristic pattern: nucleosomes are depleted directly over CTCF-bound motifs and form strongly ordered arrays in the flanking regions (solid line, Figure 4A), consistent with previous studies^47^. In contrast, unbound motifs exhibit a prominent occupancy peak with a normalized value of ∼1.25 (dashed line, Figure 4A). The *in vitro* data reveal an occupancy peak with a similar height for both bound and unbound motifs (Figure 4B). Together, these observations show that CTCF binding leads to local nucleosome depletion, whereas in the absence of CTCF, a nucleosome is intrinsically favored at the motif.

**Figure 4.**
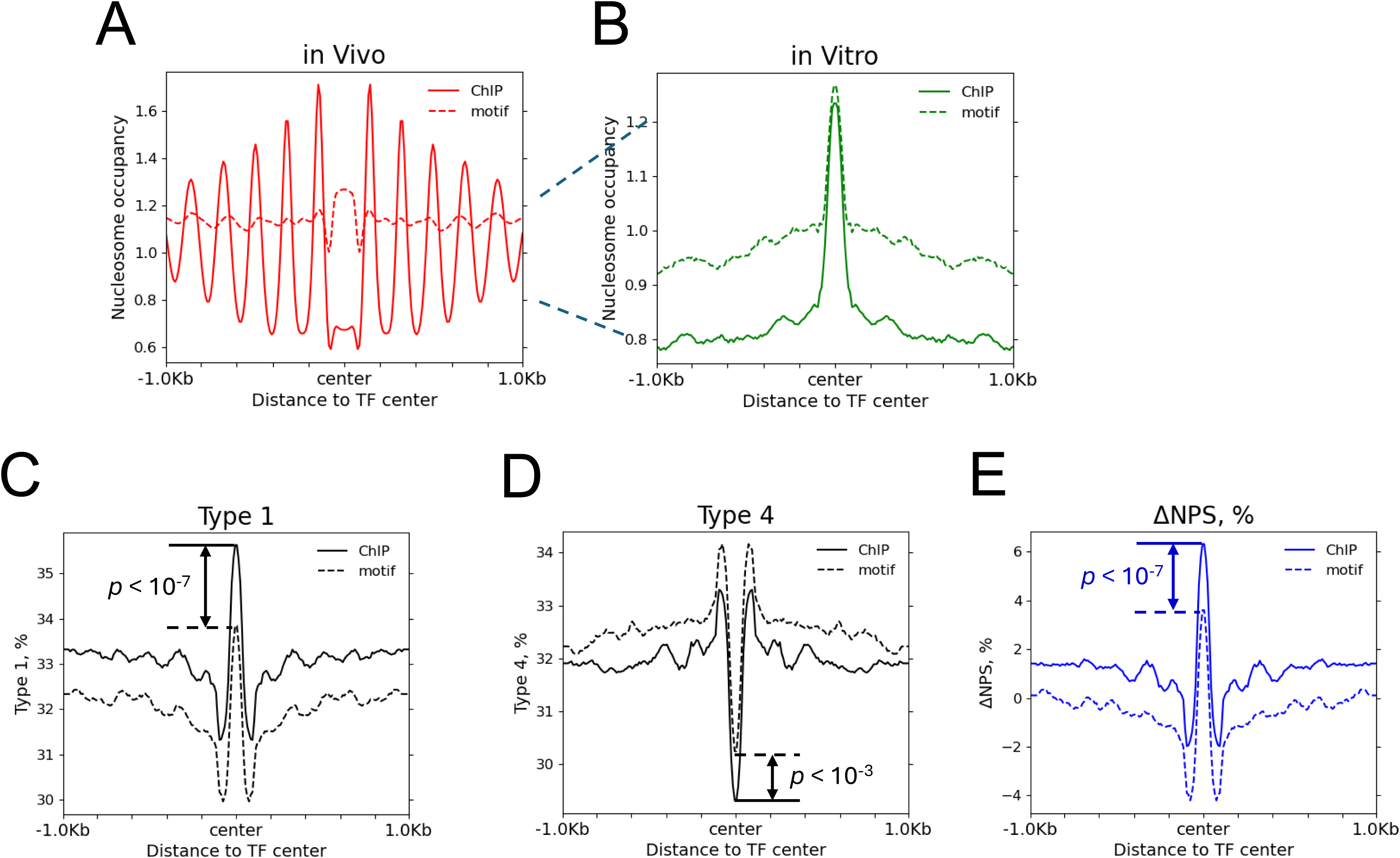
Differences in nucleosome occupancy and ΔNPS between CTCF-bound and unbound motifs. (A–B) *In vivo* (A) and *in vitro* (B) nucleosome occupancy profiles for CTCF-bound (solid lines) and unbound (dashed lines) motifs. Nucleosome occupancy at each nucleotide was normalized by dividing the number of nucleosome sequences covering that position by the genome-wide average number of nucleosome sequences per base pair. The dynamic range of *in vivo* occupancy (0.6-1.6) is substantially larger than that of *in vitro* occupancy. Dashed vertical guides are provided to facilitate comparison between profiles. (C-D) Fractions of Type-1 (C) and Type-4 (D) nucleosomes surrounding CTCF-bound (solid lines) and unbound (dashed lines) motifs. Fractions were computed in 10-bp bins across the motif-centered window. (E) ΔNPS profiles for CTCF-bound (solid line) and unbound (dashed line) motifs. Student’s *t*-tests were used to compare mean nucleosome-type fractions (C-D) and ΔNPS values (E) between bound and unbound motif sets.

Next, we examined the distribution of the four nucleosome types around bound CTCF motifs *in vivo*. Type-1 and Type-4 nucleosomes exhibit substantial changes across the motif relative to surrounding regions (Supplementary Figure S6A), whereas changes for Type-2 and Type-3 nucleosomes are minimal (Supplementary Figure S6B). This suggests that Type-1 and Type-4 nucleosomes play a dominant role in shaping nucleosome organization near CTCF-bound sites.

To investigate this further, we quantified the fractions of Type-1 and Type-4 nucleosomes near CTCF motifs. For both bound and unbound motifs, Type-1 nucleosomes are enriched at the motif center (Figure 4C). This enrichment is significantly greater for bound motifs than for unbound motifs (Student’s *t*-test, *p* < 10⁻⁷). Conversely, Type-4 nucleosomes are depleted at the motifs, with significantly stronger depletion for bound than unbound motifs (Figure 4D; Student’s *t*-test, *p* < 10⁻³).

To capture the contrasting behaviors of Type-1 and Type-4 nucleosomes, we used a previously defined metric, ΔNPS^42^, which measures the percentage difference between Type-1 and Type-4 nucleosomes in a given region. Because these two nucleosome types correspond to complementary DNA sequence patterns, WW/SS and anti-WW/SS, ΔNPS reflects the intrinsic strength of DNA-encoded nucleosome positioning. Higher ΔNPS values indicate stronger intrinsic nucleosome formation. Consistent with this, CTCF-bound motifs exhibit significantly higher ΔNPS values than unbound motifs (Figure 4D; Student’s *t*-test, *p* < 10⁻⁷), suggesting that intrinsically strong nucleosomes tend to occupy CTCF motifs and may compete with CTCF for binding.

### In vivo nucleosome occupancy and ΔNPS exhibit distinct patterns around TF binding sites

To investigate how TFs influence nucleosome organization at their genomic binding sites, we compared, for each TF, its *in vivo* NO profile with its ΔNPS profile. For CTCF, we observed a clear DIP/PEAK pattern: *in vivo* NO shows a pronounced dip over bound motifs, while the ΔNPS profile shows a corresponding peak (red and blue lines in Supplementary Figure S7A).

Plotting the *in vitro* NO profile revealed a strong peak centered on the motifs, closely resembling the ΔNPS profile (green line in Supplementary Figure S7A). This concordance reinforces the conclusion that ΔNPS captures the intrinsic strength of nucleosome positioning around TF-bound motifs. Because GC content has previously been linked to *in vitro* nucleosome occupancy at TF sites^48^, we also examined GC content around CTCF motifs. Indeed, GC content peaks at the motif center, mirroring the *in vitro* NO profile (Supplementary Figure S7C).

A second pattern, DIP/DIP, was observed for SPI1: both *in vivo* NO and ΔNPS exhibited a dip at bound motifs (Supplementary Figure S7B). The nearest ΔNPS peak is located ∼70–80 bp away from the center of the motifs. The *in vitro* NO profile also shows a dip, correlating well with ΔNPS. However, GC content behaves differently here: it shows a peak at the motif (Supplementary Figure S7D), indicating that GC content does not correlate with intrinsic nucleosome occupancy in this case.

Two additional patterns were identified in other TFs. NFIC displayed a PEAK/PEAK pattern, where both *in vivo* NO and ΔNPS profiles show peaks at bound motifs (Supplementary Figure S8A). NFYA showed a PEAK/DIP pattern, with *in vivo* NO showing a peak but ΔNPS showing a dip (Supplementary Figure S8B). For both TFs, the *in vitro* NO profile matches the ΔNPS profile. GC content correlates with intrinsic NO for NFIC (Supplementary Figure S8C), but not for NFYA (Supplementary Figure S8D).

Using these patterns, we classified 515 mammalian TFs and 232 yeast TFs into four groups, DIP/PEAK, DIP/DIP, PEAK/PEAK, and PEAK/DIP, according to their *in vivo* NO and ΔNPS profiles (Supplementary Tables S7-S9). Representative average profiles for each group are shown in Figure 5A-D. We also identified an AMBIGUOUS category for TFs lacking either a clear peak or dip in their *in vivo* NO profiles (Supplementary Figure S9).

**Figure 5.**
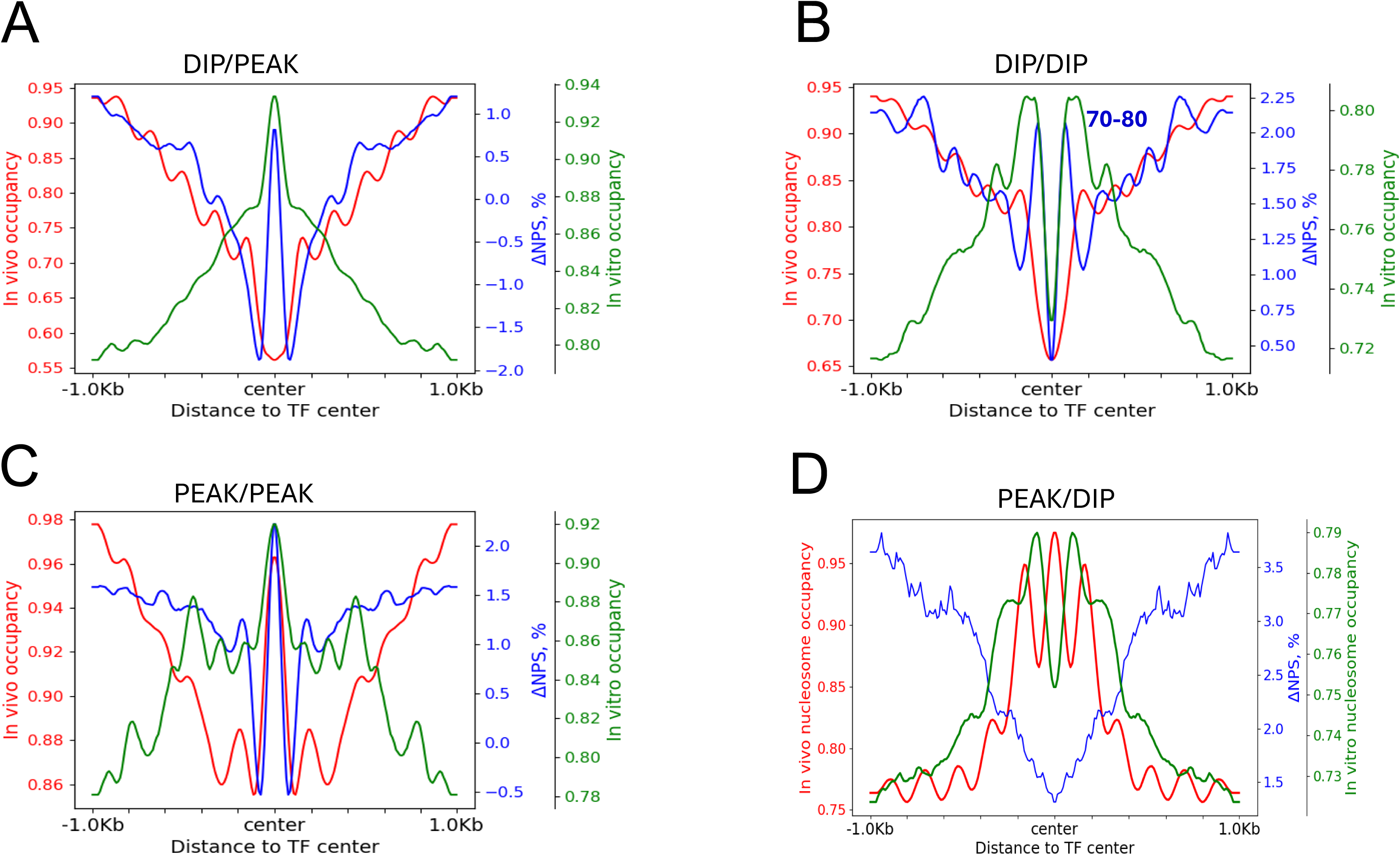
Representative nucleosome occupancy and ΔNPS profiles for the four TF pattern groups. Average profiles of *in vivo* nucleosome occupancy (red), *in vitro* nucleosome occupancy (green), and ΔNPS (blue) are shown for TFs classified into the DIP/PEAK (A), DIP/DIP (B), PEAK/PEAK (C), and PEAK/DIP (D) groups. For GM12878 cells, the DIP/PEAK, DIP/DIP, and PEAK/PEAK groups include 70, 39, and 14 TFs, respectively. Because GM12878 contains no TFs in the PEAK/DIP group, 15 TFs from HeLa cells belonging to this category were used for panel (D). All profiles are symmetrized relative to the motif center (position 0). In the DIP/DIP group (B), the ΔNPS peak closest to the motif occurs approximately 70–80 bp from the center.

Because about 20% of the 515 mammalian TFs exhibit different profile patterns across cell lines (Supplementary Table S7), we classified TFs into two datasets: a stringent set containing 413 TFs and a non-stringent set containing 515 TFs (Supplementary Table S10). The stringent set includes only TFs with a single dominant profile pattern across all examined cell lines. In contrast, the non-stringent set includes all TFs, including those with tied dominant patterns, which are assigned to each corresponding profile group.

Importantly, these profile groups found in human TFs are conserved across species: the same categories are observed in mouse TFs (Supplementary Figure S10) and yeast TFs (Supplementary Figure S11). This indicates that these patterns likely represent evolutionarily conserved modes of nucleosome-TF interactions.

## Discussion

Based on our data, we propose a unified model for the modes of nucleosome-TF interactions *in vivo* (Figure 6) using the stringent set (Supplementary Table S10).

1. DIP/PEAK group (41% of TFs). These TFs have strong intrinsic nucleosomes located directly over their binding motifs. The *in vivo* depletion of nucleosomes indicates direct competition between TFs and these intrinsically positioned nucleosomes. In other words, TF binding displaces strong DNA-encoded nucleosomes.
2. DIP/DIP group (32% of TFs). These TFs have strong intrinsic nucleosomes positioned ∼70–80 bp away from the motif center. Thus, motifs are situated either in linker DNA or near the nucleosome edge, making them easily accessible. The absence of a strong DNA-encoded nucleosome at the motif itself is likely to contribute to the NDRs observed *in vivo*.
3. PEAK/PEAK group (11% of TFs). These TFs have strong intrinsic nucleosomes centered on their motifs. The enrichment of nucleosomal DNA *in vivo* suggests that these TFs bind DNA while it remains wrapped around a nucleosome, binding “at the center” relative to the nucleosome ends, though not necessarily at the dyad. These TFs appear to recognize or stabilize a well-positioned nucleosome.
4. PEAK/DIP group (3% of TFs). These TFs lack strong DNA-encoded nucleosomes near their motifs (ΔNPS shows a dip), yet nucleosomes are enriched *in vivo*. This suggests that nucleosome positioning is imposed not by DNA sequence but by protein factors such as chromatin remodelers or even the TFs themselves.

**Figure 6.**
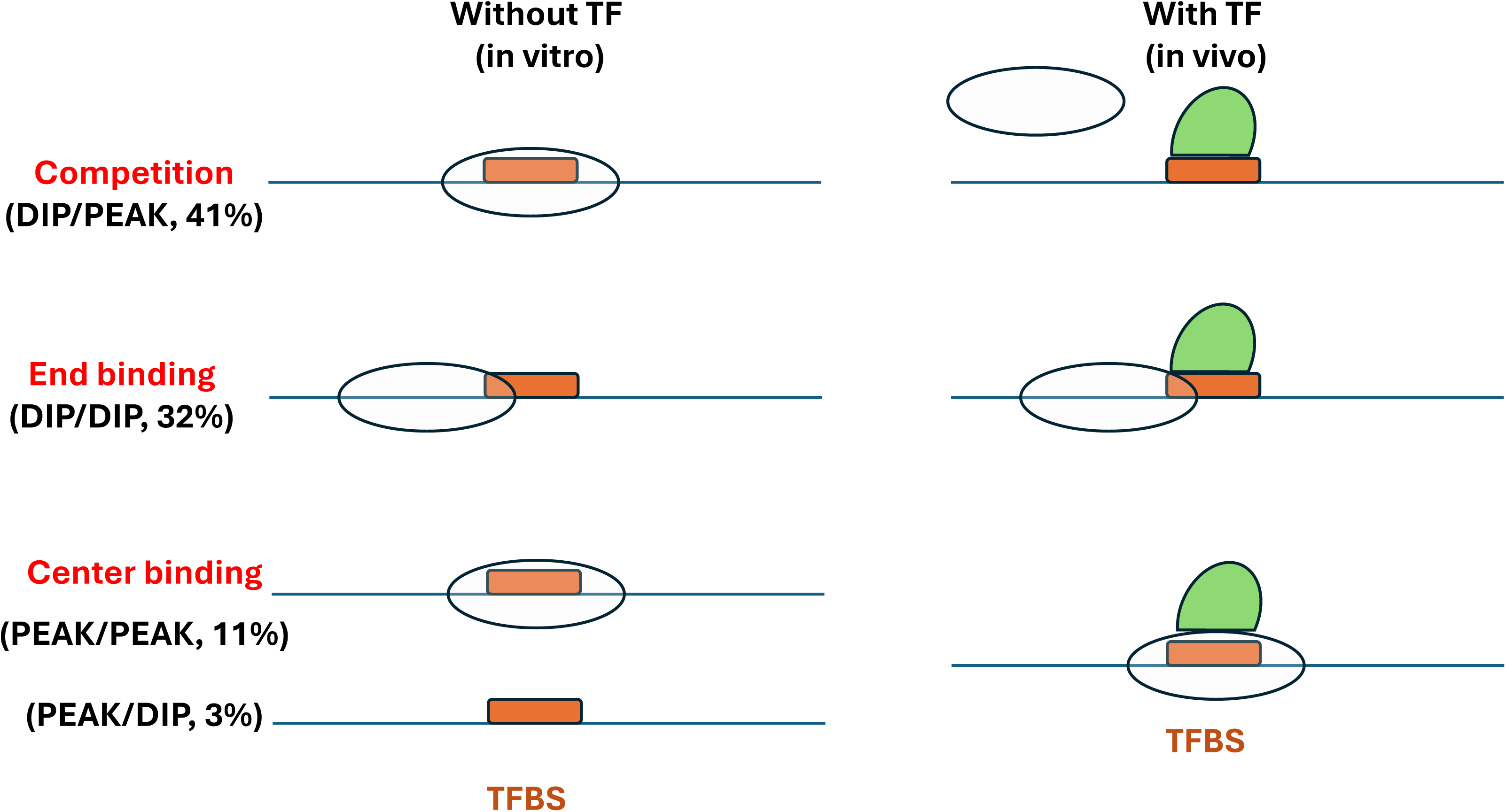
A model illustrating how DNA-encoded nucleosome patterns determine modes of nucleosome-TF interactions. Schematic representations of the four major TF profile groups, DIP/PEAK, DIP/DIP, PEAK/PEAK, and PEAK/DIP, highlight how intrinsic DNA sequence patterns shape nucleosome positioning and influence transcription factor binding. Percentages reflect the proportion of TFs in the stringent dataset. The AMBIGUOUS group (13% of TFs) is not shown because it lacks a clear peak or dip in *in vivo* nucleosome occupancy at motif centers. Their modes of nucleosome interaction appear heterogeneous or context-dependent and require further investigation.

Our data indicates that, for DIP/DIP TFs, NDRs may arise because the local DNA sequence disfavors the formation of strong, stable nucleosomes. In this case, the genomic region is intrinsically nucleosome-poor, and TF binding does not require competition with nucleosomes. In contrast, TFs in the DIP/PEAK group are associated with strong intrinsic nucleosomes positioned over their motifs, as reflected by high ΔNPS values. For these TFs, NDRs likely result from direct competition, in which DNA-encoded nucleosomes are displaced or excluded upon TF binding.

Together, these results show that NDRs can arise through at least two distinct sequence-driven mechanisms: (i) the absence of strong intrinsic nucleosomes (DIP/DIP) and (ii) TF–nucleosome competition at sites with strong intrinsic nucleosomes (DIP/PEAK).

## Online Methods

### Genome-wide nucleosome positioning (MNase-seq) datasets

Both *in vitro* and *in vivo* nucleosome maps from yeast and higher eukaryotes were used in this study, comprising ten *in vivo* datasets and two *in vitro* datasets (Supplementary Table S1). The ten *in vivo* nucleosome maps included eight human datasets, one mouse dataset, and one yeast dataset. The human datasets were derived from eight cell lines: GM12878 (GSM920558), K562 (GSM920557), MCF-7 (GSE51097), H1-hESC (GSM1194220), HepG2 (GSM3718063), HeLa (GSE100401), HCT116 (GSM3889219), and IMR90 (GSM1095279). The mouse dataset was generated from mouse embryonic stem cells (mESC; GSM2183911), and the yeast dataset was obtained from cells grown to late-log phase in synthetic complete medium (GSM651368). Two *in vitro* nucleosome positioning datasets were also analyzed—one from humans (GSE25133) and one from yeast (GSE13622).

Nucleosome occupancy maps were obtained either directly from ENCODE or generated by aligning MNase-seq short reads to the corresponding reference genomes using Bowtie2 (Supplementary Figure S12). For example, occupancy tracks (bigWig format) for GM12878 and K562 were downloaded from ENCODE. To generate occupancy profiles for each of the four nucleosome types, every nucleosome was first assigned to Type-1, Type-2, Type-3, or Type-4 based on its DNA sequence pattern. Genome-wide nucleosome occupancy for each type was then computed from dyad counts using a center-weighted scheme^49^.

For human cell lines without pre-computed occupancy files, MNase-seq reads in the FASTQ format were mapped to the genome using Bowtie2 with the following parameters: --sensitive -I 0 -X 500 --no-discordant. Mapped reads were extended from their 5′ ends to a canonical 147-bp fragment length. Nucleosome occupancy for these datasets was calculated following our previous protocols^46^.

The mESC dataset was provided as center-weight scores, which we converted to BedGraph format before downstream processing. For yeast datasets and both *in vitro* nucleosome maps, occupancy profiles were generated directly from dyad counts using the same center-weighting procedure^49^.

### Classification of nucleosomes with different NPS patterns

Nucleosomal DNA sequences were classified into four types, e.g., Type-1, Type-2, Type-3, and Type-4, based on the relative abundance of WW and SS dinucleotides in minor- and major-GBS^42^. Type-1 nucleosomes exhibit the canonical WW/SS pattern. For a 147-bp NCP fragment, if (1) the total number of WW dinucleotides in the 12 minor-GBS is greater than that in the 12 major-GBS, and (2) the total number of SS dinucleotides in the 12 major-GBS is greater than that in the 12 minor-GBS, the fragment is classified as Type-1. Type-4 nucleosomes display the anti-WW/SS pattern, the inverse of Type-1. A 147-bp fragment is assigned to Type-4 if (1) WW dinucleotides are more abundant in the 12 major-GBS than in the minor-GBS, and (2) SS dinucleotides are more abundant in the 12 minor-GBS than in the major-GBS. Type-2 and Type-3 nucleosomes represent “mixed” patterns. Type-2 fragments contain more WW and more SS dinucleotides in minor-GBS than in major-GBS. Type-3 fragments contain more WW and more SS dinucleotides in major-GBS than in minor-GBS.

For any genomic region, we quantify the relative contributions of the two complementary sequence patterns using ΔNPS, defined as ΔNPS = Type 1 (%) – Type 4 (%), which reflects the balance between WW/SS and anti-WW/SS nucleosome patterns (Supplementary Figure S13).

Type-1 through Type-4 nucleosomes were identified in both *in vivo* (GM12878; Supplementary Table S2) and *in vitro* datasets (Supplementary Table S3). Overall, the number of nucleosomes detected in the *in vitro* dataset is approximately one-third of that detected *in vivo* (compare Supplementary Tables S2 and S3).

### Genome-wide protein-DNA interaction (ChIP-seq) datasets

Human and mouse ChIP-seq datasets were obtained from ENCODE. For each TF, we downloaded the processed BED files containing mapped ChIP fragments. For approximately 50% of TFs across all analyzed cell lines, the position-specific scoring matrix of the corresponding TF binding motif was obtained by matching TF names to entries in the HOCOMOCO v11, JASPAR 2022, and FactorBook motif databases.

TF binding sites were identified by scanning each ChIP fragment with the corresponding PSSM using MEME suite tools. When multiple motif hits were present within a fragment, the highest-scoring site was retained for downstream analysis. For yeast, we used the processed ChIP-seq dataset^50^ in which the centers of binding motifs were already provided for each ChIP fragment.

For CTCF, unbound motifs were generated as follows. The human genome was scanned with the CTCF PSSM (CTCF_human_h11MO.meme) using FIMO with a significance threshold of 10⁻⁴. Motifs overlapping ChIP-identified binding sites were removed to obtain a set of unbound motifs. The same number of unbound motifs as their ChIP-derived counterparts were then randomly selected for further analysis.

### Classification of TF patterns using in vivo nucleosome occupancy and ΔNPS profiles

Transcription factors were classified into five categories, DIP/PEAK, DIP/DIP, PEAK/PEAK, PEAK/DIP, and AMBIGUOUS, based on the patterns of *in vivo* NO and ΔNPS profiles centered on TF binding motifs (position 0 in Figure 5).

Classification criteria were defined using the relative positions of global maxima and minima within each profile, normalized by the full dynamic range of the signal:

DIP/PEAK: The *in vivo* NO profile exhibits a global minimum (or a value below the lower one-third of its dynamic range), while the ΔNPS profile exhibits a global maximum (or a value above the upper one-third of its range) at the motif center.

DIP/DIP: Both the *in vivo* NO and ΔNPS profiles show a global minimum (or values within the lowest one-third of their respective ranges) at the motif center.

PEAK/PEAK: Both the *in vivo* NO and ΔNPS profiles display a global maximum (or values within the upper one-third of their ranges) at the motif center.

PEAK/DIP: The *in vivo* NO profile shows a global maximum (upper one-third of its range), while the ΔNPS profile shows a global minimum (lower one-third of its range) at the motif center.

AMBIGUOUS: TFs whose *in vivo* NO profiles exhibit no clear peak or dip, i.e., values at the motif center fall between the lower and upper one-third thresholds, were placed in the AMBIGUOUS group.

This classification scheme ensures that each TF is assigned based on robust, quantitative features of nucleosome organization and intrinsic DNA-encoded nucleosome strength around its binding motifs.

### Pattern Labeling

Some transcription factors exhibited more than one TF pattern across different cell lines. To derive a consensus label for each TF, we counted the occurrences of all patterns observed for that TF and assigned as its consensus classification the pattern(s) tied for the highest frequency. Thus, a TF could have a single consensus label (e.g., DIP/DIP) or multiple labels if ties occurred (e.g., DIP/DIP, DIP/PEAK, and AMBIGUOUS). In addition to this comprehensive dataset, we generated a stringent dataset in which TFs with more than one consensus label were removed. The stringent dataset, therefore, contains only TFs that exhibit a single, consistent pattern across all examined cell lines.

### Inter-nucleosome distance correlation function

To quantify the relative spatial organization of nucleosomes, we applied DAC and DCC functions^46^. For every pair of nucleosomal DNA fragments, we calculated the distance between their start positions. When multiple identical nucleosome sequences occurred at the same genomic location, all occurrences were counted multiplicatively—for example, if two specific sequences occurred 5 and 10 times, respectively, the DAC contribution for that pair was counted as 5 × 10 = 50.

## Acknowledgments

The authors thank Victor B. Zhurkin for his guidance and stimulating discussions. We are grateful to the Thomas H. Gosnell School of Life Sciences and the College of Science at RIT for administrative and financial support. We also acknowledge the RIT Research Computing team for providing computational resources and support.

## Availability of methods

All computational methods and pipelines are available in GitHub https://github.com/rit-cui-lab/Hidden-DNA-codes-influence-the-interaction-between-nucleosomes-and-transcription-factors/tree/main

## References

1. Kornberg, R.D. & Lorch, Y. Twenty-five years of the nucleosome, fundamental particle of the eukaryote chromosome. Cell 98, 285–294 (1999).

2. Jiang, C. & Pugh, B.F. Nucleosome positioning and gene regulation: advances through genomics. Nat. Rev. Genet. 10, 161–172 (2009).

3. Struhl, K. & Segal, E. Determinants of nucleosome positioning. Nat. Struct. Mol. Biol. 20, 267–273 (2013).

4. Blomquist, P., Li, Q. & Wrange, O. The affinity of nuclear factor for its DNA site is drastically reduced by nucleosome organization irrespective of its rotational or translational position. J. Biol. Chem. 27, 153–159 (1996).

5. Owen-Hughes, T. & Workman, J.L. Experimental analysis of chromatin function in transcription control. Crit. Rev. Eukaryot. Gene Exp. 4, 403–441 (1994).

6. Wang, J., Zhuang, J., Iyer, S., Lin, X., Whitfield, T.W., Greven, M.C., Pierce, B.G., Dong, X., Kundaje, A., Cheng, Y., Rando, O.J., Birney, E., Myers, R.M., Noble, W.S., Snyder, M. and Weng, Z. Sequence features and chromatin structure around the genomic regions bound by 119 human transcription factors. Genome Res. 22, 1798–1812 (2012).

7. Iwafuchi-Doi, M. & Zaret, K.S. Pioneer transcription factors in cell reprogramming. Genes Dev. 28, 2679–2692 (2014).

8. Zaret, K.S. & Carroll, J.S. Pioneer transcription factors: establishing competence for gene expression. Genes Dev. 25, 2227–2241 (2011).

9. Cirillo, L.A. & Zaret, K.S. An early developmental transcription factor complex that is more stable on nucleosome core particles than on free DNA. Mol. Cell 4, 961–969 (1999).

10. Sekiya, T., Muthurajan, U.M., Luger, K., Tulin, A.V. & Zaret, K.S. Nucleosome-binding affinity as a primary determinant of the nuclear mobility of the pioneer transcription factor FoxA. Genes Dev. 23, 804–809 (2009).

11. Mivelaz, M., Cao, A.M., Kubik, S., Zencir, S., Hovius, R., Boichenko, I., Stachowicz, A.M., Kurat, C.F., Shore, D., & Fierz, B. Chromatin fiber invasion and nucleosome displacement by the Rap1 transcription factor. Mol. Cell 77, 488–500 (2020).

12. Donovan, B.T., Chen, H., Jipa, C., Bai, L., & Poirier, M.G. Dissociation rate compensation mechanism for budding yeast pioneer transcription factors. Elife 8, e43008 (2019).

13. Michael, A.K., Grand, R.S., Isbel, L., Cavadini, S., Kozicka, Z., Kempf, G., Bunker, R.D., Schenk, A.D., Graff-Meyer, A., Pathare, G.R., Weiss, J., Matsumoto, S. Burger, L., Schubeler, D. & Thoma, N.H. Mechanisms of OCT4-SOX2 motif readout on nucleosomes. Science 368, 1460–1465 (2020).

14. Dodonova, S.O., Zhu, F., Dienemann, C., Taipale, J. & Cramer, P. Nucleosome-bound SOX2 and SOX11 structures elucidate pioneer factor function. Nature 580, 669–672 (2020).

15. Tanaka, H., Takizawa, Y., Takaku, M., Kato, D., Kumagawa, Y., Grimm, S.A., Wade, P.A. & Kurumizaka, H. Interaction of the pioneer transcription factor GATA3 with nucleosomes. Nat. Comm. 11, 4136 (2020).

16. Zhu, F., Farnung, L., Kaasinen, E., Sahu, B., Yin, Y., Wei, B., Dodonova, S.O., Nitta, K.R., Morgunova, E. & Taipale, M. The interaction landscape between transcription factors and the nucleosome. Nature 562, 76–81 (2018).

17. Garcia, M.F., Moore, C.D., Schulz, K.N., Alberto, O., Donague, G., Harrison, M.M., Zhu, H. & Zaret, K.S. Structural features of transcription factors associating with nucleosome binding. Mol. Cell 75, 921–932 (2019).

18. Lowary, P.T. & Widom, J. New DNA sequence rules for high affinity binding to histone octamer and sequence-directed nucleosome positioning. J. Mol. Biol. 276, 19–42 (1998).

19. Mivelaz, M., Cao, A-M., Kubik, S., Zencir, S., Hovius, R., Boichenko, I., Stachowicz, A.M., Kurat, C.F., Shore, D. & Fierz, B. Chromatin fiber invasion and nucleosome displacement by the Rap1 transcription factor. Mol. Cell 77, 488–500 (2020).

20. Laptenko, O., Beckerman, R., Freulich, E. & Prives, C. p53 binding to nucleosomes within the p21 promoter in vivo leads to nucleosome loss and transcriptional activation. Proc. Natl. Acad. Sci. U.S.A. 108, 10385–10390 (2011).

21. Soufi, A., Garcia, M.F., Jaroszewicz, A., Osman, N., Pellegrini, M. & Zaret, K.S. Pioneer transcription factors target partial DNA motifs on nucleosomes to initiate reprogramming. Cell 161, 555–568 (2015).

22. Li, Q. & Wrange, O. Accessibility of a glucocorticoid response element in a nucleosome depends on its rotational positioning. Mol. Cell Biol. 15, 4375–4384 (1995).

23. Sahu, G., Wang, D., Chen, C.B., Zhurkin, V.B., Harrington, R.E., Appella, E., Hager, G.L. & Nagaich, A.K. p53 binding to nucleosomal DNA depends on the rotational positioning of DNA response element. J. Biol. Chem. 285, 1321–1332 (2010).

24. Cui, F. & Zhurkin, V.B. Rotational positioning of nucleosomes facilitates selective binding of p53 to response elements associated with cell cycle arrest. Nucleic Acids Res. 42, 836–847 (2014).

25. Li, S., Zheng, E.B., Zhao, L. & Liu, S. Nonreciprocal and conditional cooperativity directs the pioneer activity of pluripotency transcription factors. Cell Rep. 28, 2689–2703 (2019).

26. Lu, Q., Wallrath, L.L. & Elgin, S.C. Nucleosome positioning and gene regulation. J. Cell. Biochem. 55, 83–92 (1994).

27. Mengeritsky, G. & Trifonov, E.N. Nucleotide sequence-directed mapping of the nucleosomes. Nucleic Acids Res. 11, 3833–3851 (1983).

28. Zhurkin, V.B. Specific alignment of nucleosomes on DNA correlates with periodic distribution of purine-pyrimidine and pyrimidine-purine dimers. FEBS Lett. 158, 293–297 (1983).

29. Uberbacher, E.C., Harp, J.M. & Bunick, G.J. DNA sequence patterns in precisely positioned nucleosome. J. Biomol. Struct. Dyn. 6, 105–120 (1988).

30. Ioshikhes, I., Bolshoy, A. & Trifonov, E.N. Preferred positions of AA and TT dinucleotides in aligned nucleosome DNA sequences. J. Biomol. Struct. Dyn. 9, 1111–1117 (1992).

31. Baldi, P., Brunak, S., Chauvin, Y. & Krogh, A. Naturally occurring nucleosome positioning signals in human exons and introns. J. Mol. Biol. 263, 503–510 (1996).

32. Ioshikhes, I., Bolshoy, A., Derenshteyn, K., Borodovsky, M. & Trifonov, E.N. Nucleosome DNA sequence pattern revealed by multiple alignment of experimentally mapped sequences. J. Mol. Biol. 262, 129–139 (1996).

33. Kogan, S.B., Kato, M., Kiyama, R. & Trifonov, E.N. (2006) Sequence structure of human nucleosome DNA. J. Biomol. Struct. Dyn. 24, 43–48 (2006).

34. Satchwell, S.C., Drew, H.R. & Travers, A.A. Sequence periodicities in chicken nucleosome core DNA. J. Mol. Biol. 191, 659–675 (1986).

35. Muthurajan, U.M., Bao, Y., Forsberg, L.J., Edayathumangalam, R.K., Suto, R.K., Chakravarthy, S., Dyer, P.N. & Luger, K. Structure and dynamics of nucleosomal DNA. Biopolymers 68, 547–556 (2003).

36. Sullivan, S.A. & Landsman, D. Characterization of sequence variability in nucleosome core histone folds. Proteins 52, 454–465 (2003).

37. Davey, C.A., Sargent, D.F., Luger, K., Maeder, A.W. & Richmond, T.J. Solvent mediated interactions in the structure of the nucleosome core particle at 1.9Å resolution. J. Mol. Biol. 391, 1097–1113 (2002).

38. Rohs, R., West, S.M., Sosinsky, A., Liu, P., Mann, R.S. & Honig, B. (2009) The role of DNA shape in protein-DNA recognition. Nature 461, 1248–1253 (2009).

39. Wang, D., Ulyanov, N.B. & Zhurkin, V.B. Sequence-dependent Kink-and-Slide deformations of nucleosomal DNA facilitated by histone arginines bound in the minor groove. J. Biomol. Struct. Dyn. 27, 843–859 (2010).

40. West, S.M., Rohs, R., Mann, R.S. & Honig, B. Electrostatic interactions between arginines and the minor groove in the nucleosomes. J. Biomol. Struct. Dyn. 27, 861–866 (2010).

41. Ioshikhes, I., Hosid, S. & Pugh, B.F. Variety of genomic DNA patterns for nucleosome positioning. Genome Res. 21, 1863–1871 (2011).

42. Wright, G.M. & Cui, F. The nucleosome position-encoding WW/SS sequence pattern is depleted in mammalian genes relative to other eukaryotes. Nucleic Acids Res. 47, 7942–7954 (2019).

43. Nikitina, T., Guiblet, W.M., Cui, F. & Zhurkin, V.B. Histone N-tails modulate sequence-specific positioning of nucleosomes. J. Biol. Chem. 301, 108138 (2025).

44. Valouev, A., Johnson, S.M., Boyd, S.D., Smith, C.L., Fire, A.Z. & Sidow, A. Determination of nucleosome organization in primary human cells. Nature 474, 516–520 (2011).

45. Kundaje, A., Kyriazopoulou-Panagiotopoulou, S., Libbrecht, M., Smith, C.L., Raha, D., Winters, E.E., Johnson, S.M., Snyder, M., Batzoglou, S. & Sidow, A. Ubiquitous heterogeneity and asymmetry of the chromatin environment at regulatory elements. Genome Res. 22, 1735–1747 (2012).

46. Cui, F., Cole, H.A., Clark, D.J. & Zhurkin, V.B. Transcriptional activation of yeast genes disrupts intragenic nucleosome phasing. Nucleic Acids Res. 40, 10753–10764 (2012).

47. Teif, V.B., Vainshtein, Y., Caudron-Herger, M., Mallm, J.-P., Marth, C., Hofer, T. & Rippe, K. Genome-wide nucleosome positioning during embryonic stem cell development. Nat. Struct. Mol. Biol. 19, 1185–1192 (2012).

48. Tillo, D., Kaplan, N., Moore, I.K., Fondufe-Mittendorf, Y., Gossett, A.J., Field, Y., Lieb, J.D., Widom, J., Segal, E. & Hughes, T.R. High nucleosome occupancy is encoded at human regulatory sequences. PLoS One 5, e9129 (2010).

49. Voong, L.N., Xi, L., Sebeson, A.C., Xiong, B., Wang, J.P. & Wang, X. Insights into nucleosome organization in mouse embryonic stem cells through chemical mapping. Cell 167, 1555–1570 (2016).

50. Rossi, M.J., Kuntala, P.K., Lai, W.K.M., Yamada, N., Badjatia, N., Mittal, C., Kuzu, G., Bocklund, K., Farrell, N.P., Blanda, T.R., Mairose, J.D., Basting, A.V., Mistretta, K.S., Rocco, D.J., Perkinson, E.S., Kellogg, G.D., Mahony, S. & Pugh, B.F. A high-resolution protein architecture of the budding yeast genome. Nature 592, 309–314 (2021).

