## Supplementary Figures for "Intrinsic DNA codes govern distinct modes of nucleosome-transcription factor interactions"

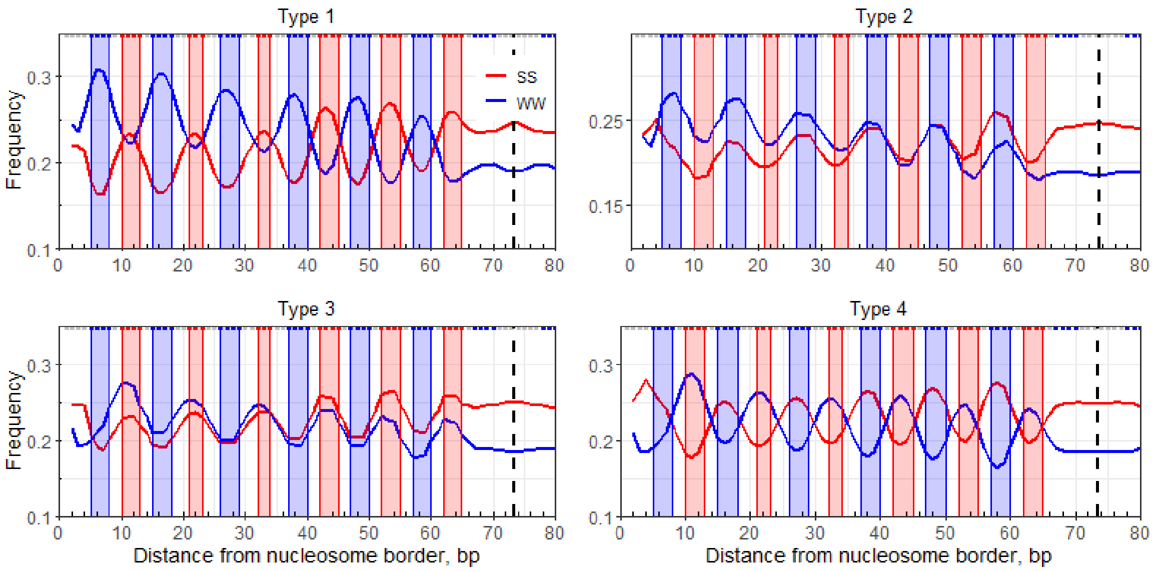

**Figure S1. Four nucleosomal DNA sequence patterns identified in the human *in vitro* dataset.** Frequencies of AA, TT, AT, and TA dinucleotides (WW, shown in blue) and GG, CC, GC, and CG dinucleotides (SS, shown in red) are plotted across nucleosomal DNA and symmetrized with respect to the dyad (dashed line). Three–base pair running averages of WW and SS frequencies are shown for the four nucleosome types: Type-1, Type-2, Type-3, and Type-4. Blue and red shaded regions indicate the positions of minor- and major-groove bending sites (minor-GBS and major-GBS), respectively. Type-1 nucleosomes correspond to the canonical WW/SS pattern, whereas Type-4 nucleosomes correspond to the inverse anti-WW/SS pattern.

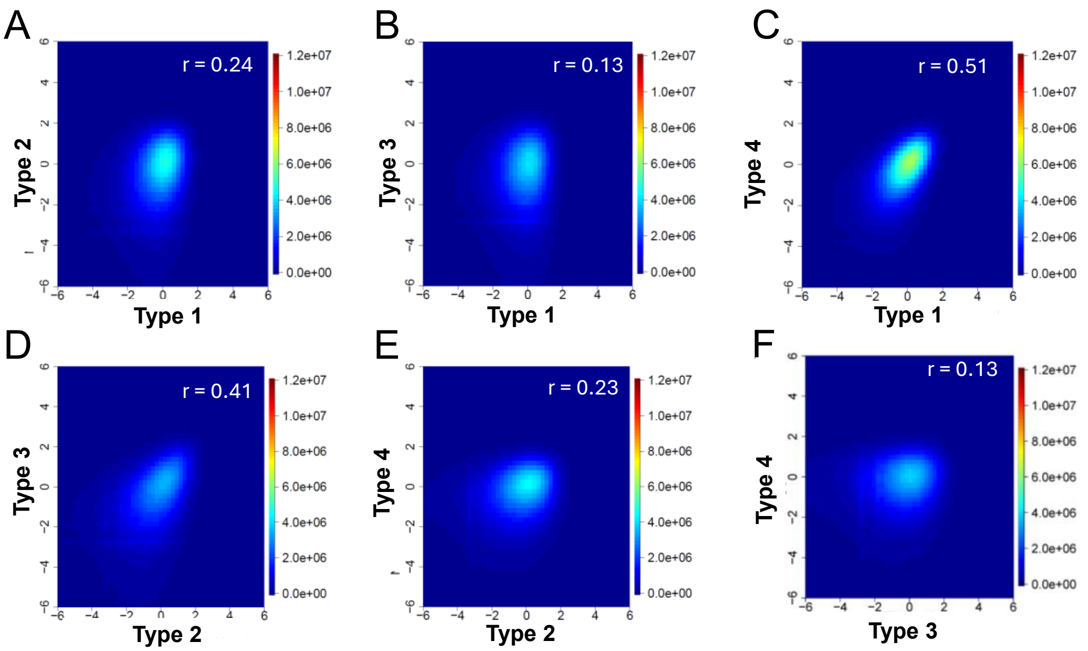

**Figure S2. Comparison of genome-wide *in vivo* nucleosome occupancy across nucleosome types.** Density scatter plots show normalized nucleosome occupancy per base pair for pairwise comparisons among nucleosome types: Type-1 vs. Type-2 (A), Type-1 vs. Type-3 (B), Type-1 vs. Type-4 (C), Type-2 vs. Type-3 (D), Type-2 vs. Type-4 (E), and Type-3 vs. Type-4 (F). Plot notation and occupancy normalization follow those described in Figure 2.

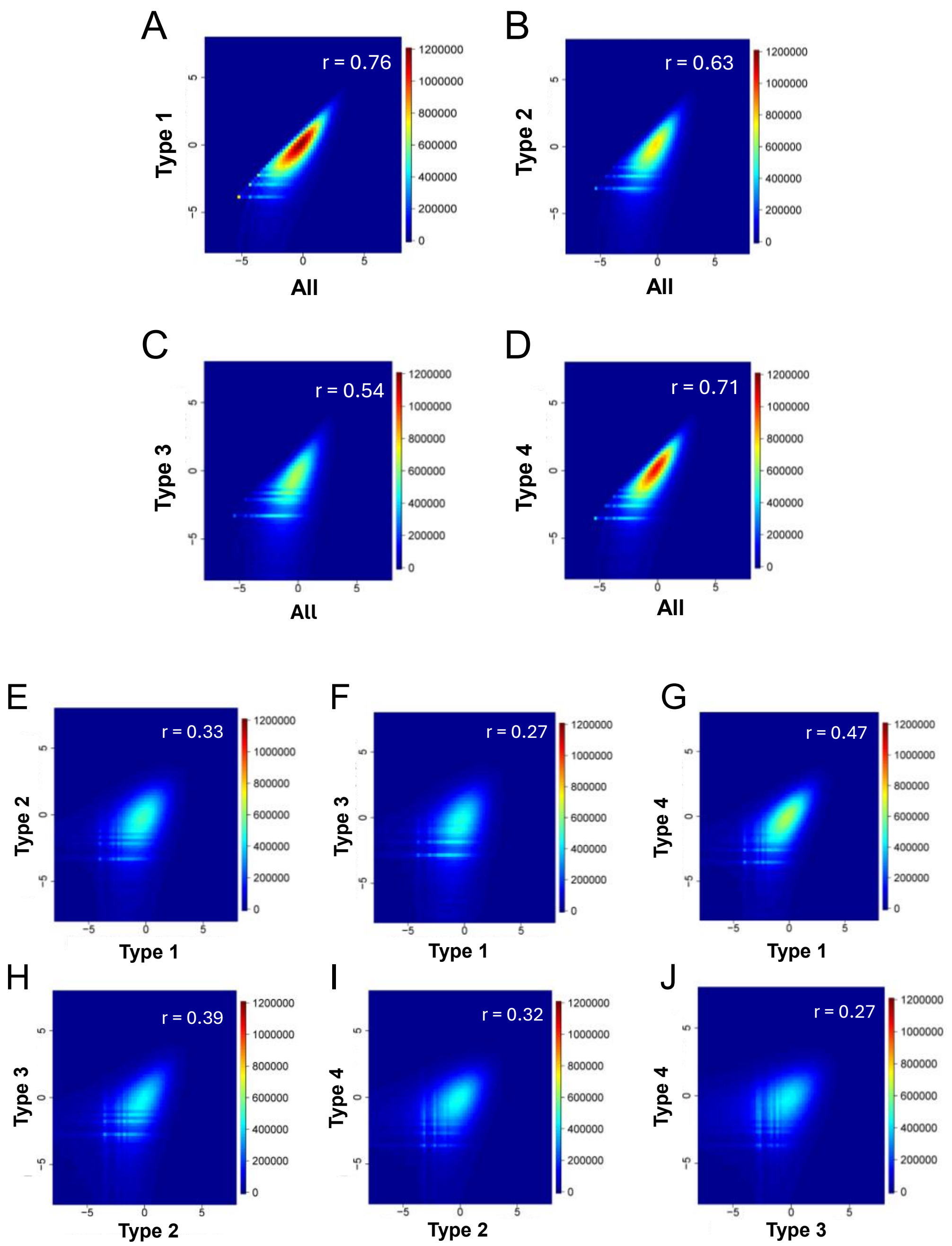

**Figure S3. Comparison of genome-wide *in vitro* nucleosome occupancy across nucleosome types.** Density scatter plots show normalized nucleosome occupancy per base pair for the complete nucleosome set compared with Type-1 (A), Type-2 (B), Type-3 (C), and Type-4 (D) nucleosome sets. Additional pairwise comparisons among nucleosome types are shown for Type-1 vs. Type-2 (E), Type-1 vs. Type-3 (F), Type-1 vs. Type-4 (G), Type-2 vs. Type-3 (H), Type-2 vs. Type-4 (I), and Type-3 vs. Type-4 (J). Plot notation and occupancy normalization methods follow those described in Figure 2.

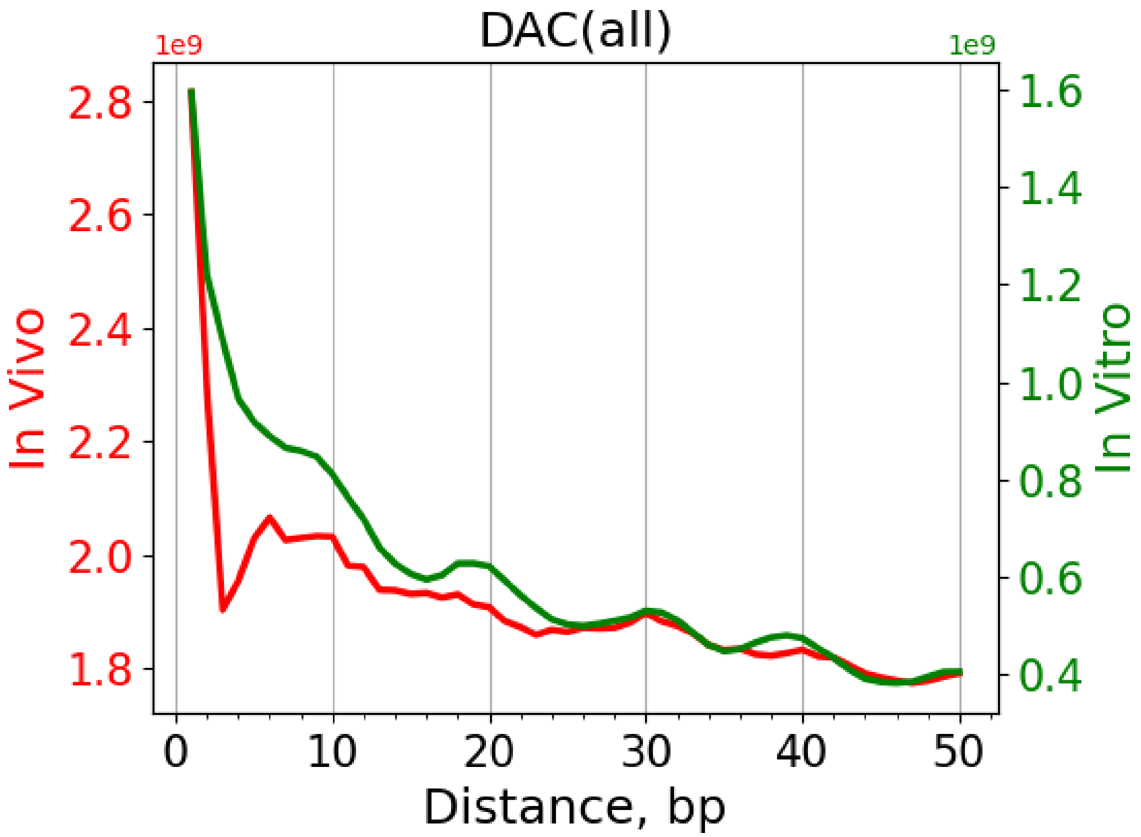

**Figure S4. Distance auto-correlation (DAC) functions for all nucleosomes in the human genome.** DAC functions were calculated genome-wide for all *in vivo* (red) and *in vitro* (green) nucleosomes. These functions reflect rotational periodicity across the full nucleosome population.

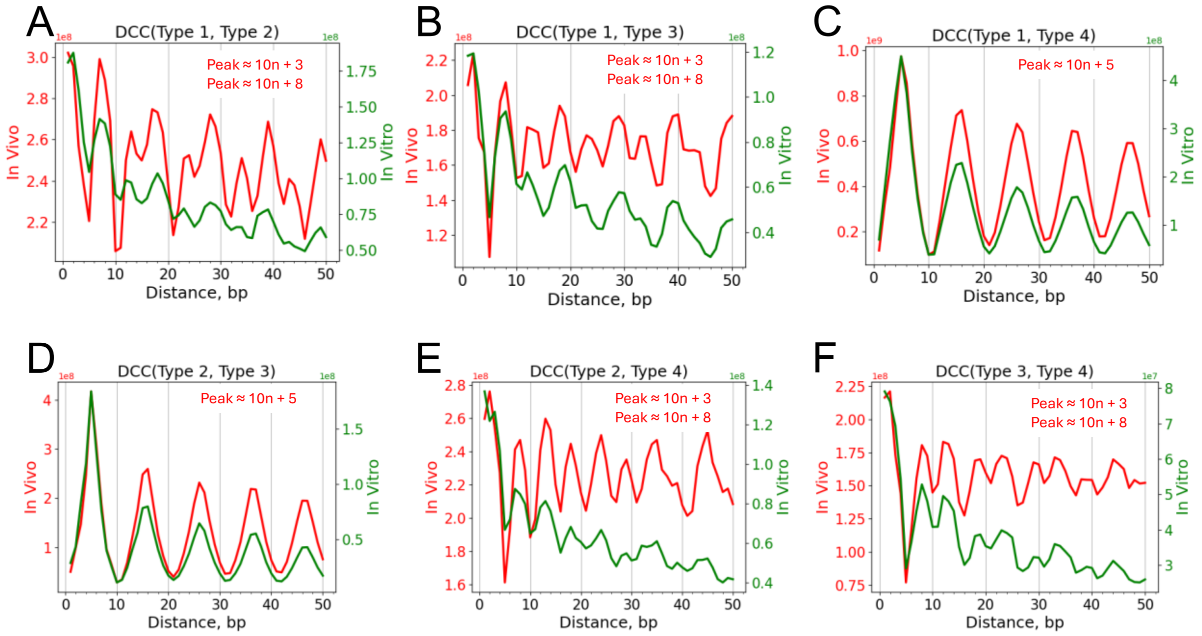

**Figure S5. Distance cross-correlation (DCC) functions between nucleosome types.** DCC functions were computed for all pairwise combinations of nucleosome types: Type-1 vs. Type-2 (A), Type-1 vs. Type-3 (B), Type-1 vs. Type-4 (C), Type-2 vs. Type-3 (D), Type-2 vs. Type-4 (E), and Type-3 vs. Type-4 (F). Both *in vivo* (red) and *in vitro* (green) nucleosome maps were used for the calculations. Peak positions for the *in vivo* DCC functions were estimated to determine the characteristic rotational offsets between nucleosome types.

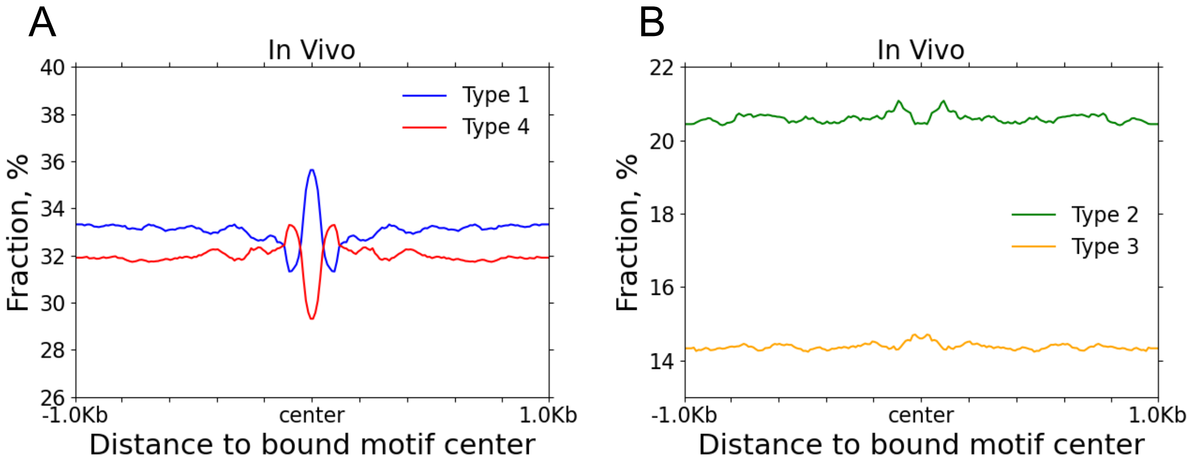

**Figure S6. Fractions of Type-1 to Type-4 *in vivo* nucleosomes around CTCF-bound motifs.** (A) Fractions of Type-1 and Type-4 nucleosomes surrounding CTCF-bound motifs. (B) Fractions of Type-2 and Type-3 nucleosomes in the same regions. Fractions were computed in 10-bp bins relative to the motif center (position 0).

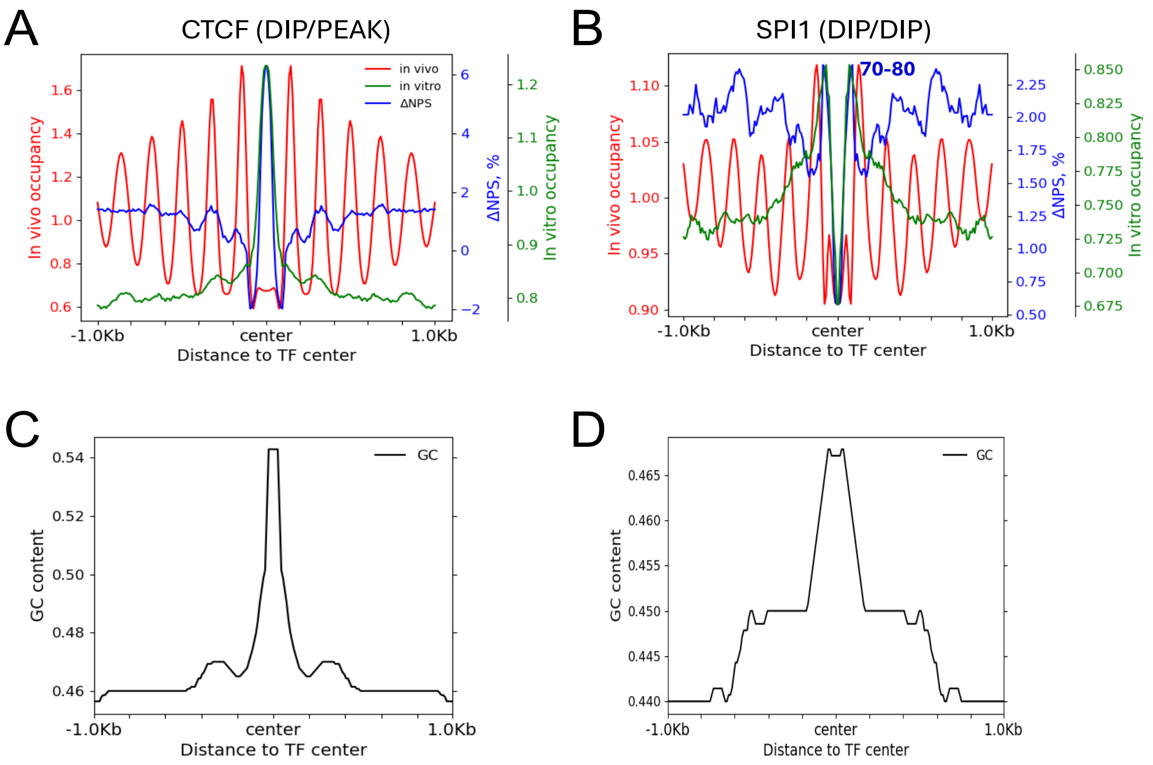

**Figure S7. Nucleosome occupancy, ΔNPS, and GC content profiles around CTCF and SPI1 binding motifs.**

(A–B) *In vivo* nucleosome occupancy (red), *in vitro* nucleosome occupancy (green), and ΔNPS values (blue) centered on CTCF (A) and SPI1 (B) bound motifs (position 0). For CTCF, the motif center coincides with a minimum in the in vivo nucleosome occupancy profile and a maximum in the ΔNPS profile, corresponding to the DIP/PEAK pattern. For SPI1, the motif center corresponds to minima in both *in vivo* occupancy and ΔNPS profiles, representing the DIP/DIP pattern. Notably, for SPI1 (B), the ΔNPS peak nearest to the motif is located ~70–80 bp away from the center. (C–D) GC content profiles around CTCF (C) and SPI1 (D) bound motifs. ΔNPS and GC content values were computed in 10-bp bins, and all profiles were symmetrized relative to the motif center (position 0).

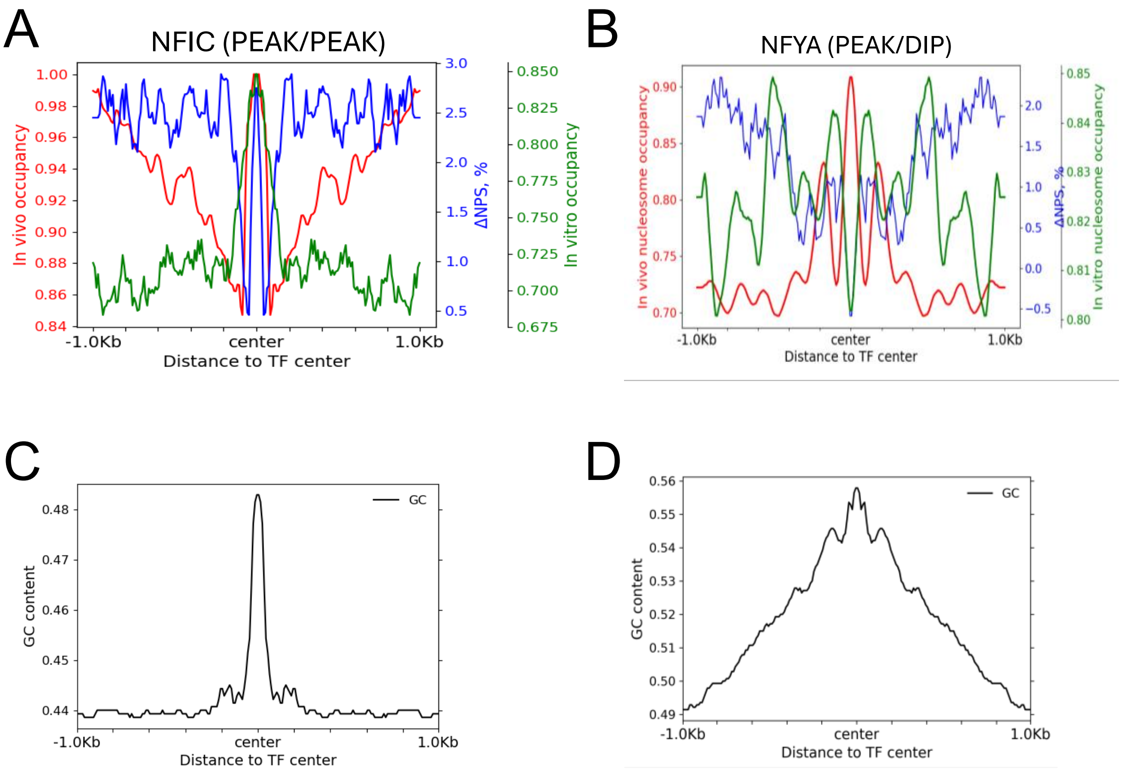

**Figure S8. Nucleosome occupancy, ΔNPS, and GC content profiles for NFIC and NFYA binding motifs.** (A–B) *In vivo* nucleosome occupancy (red), *in vitro* nucleosome occupancy (green), and ΔNPS values (blue) centered on bound motifs of NFIC in GM12878 cells (A) and NFYA in HeLa cells (B). For NFIC, the motif center corresponds to a maximum in both *in vivo* occupancy and ΔNPS, consistent with the PEAK/PEAK pattern. For NFYA, the motif center shows a maximum in *in vivo* occupancy but a minimum in ΔNPS, corresponding to the PEAK/DIP pattern. (C–D) GC content profiles surrounding NFIC (C) and NFYA (D) bound motifs. ΔNPS and GC content values were computed in 10-bp bins, and all profiles were symmetrized relative to the motif center (position 0).

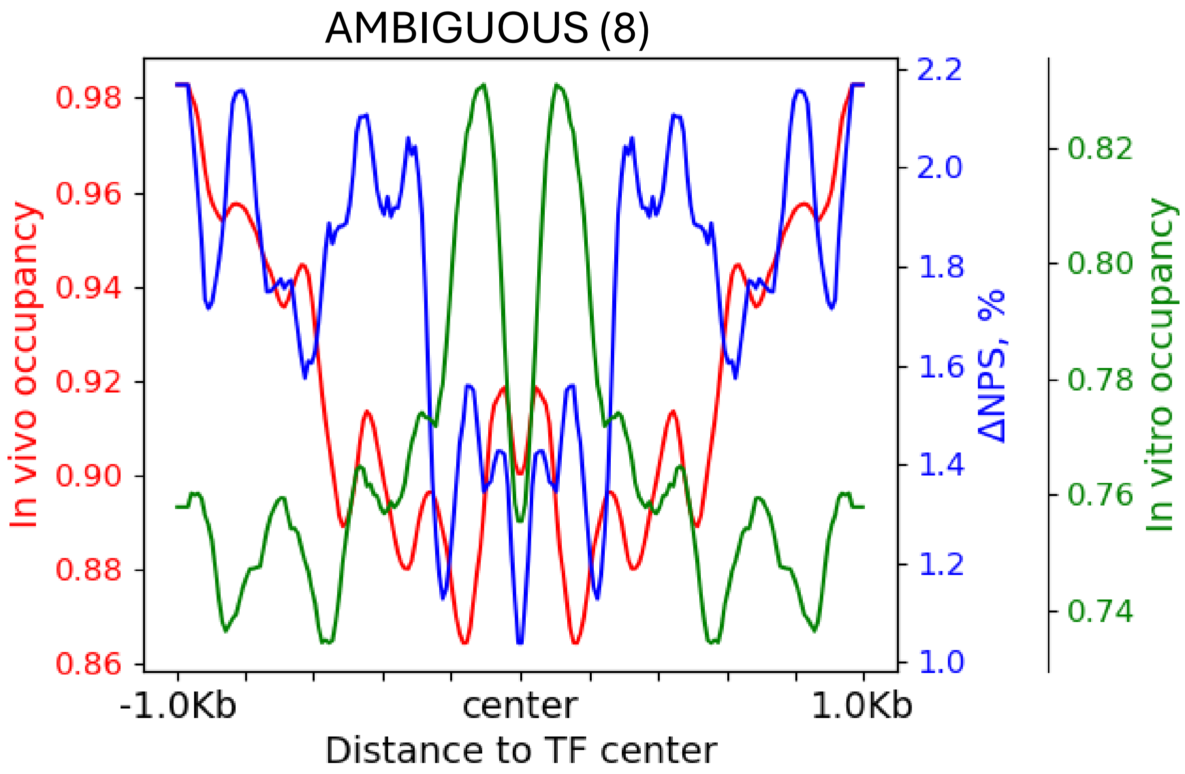

**Figure S9. Nucleosome occupancy and ΔNPS profiles for the AMBIGUOUS transcription factor group.** This group includes 8 TFs from GM12878 cells. Average profiles of *in vivo* nucleosome occupancy (red), *in vitro* nucleosome occupancy (green), and ΔNPS (blue) were calculated and plotted. All profiles were symmetrized relative to the motif center (position 0). In this group, the *in vivo* nucleosome occupancy at the motif center does not correspond to either a clear maximum or a minimum, defining the profile as AMBIGUOUS.

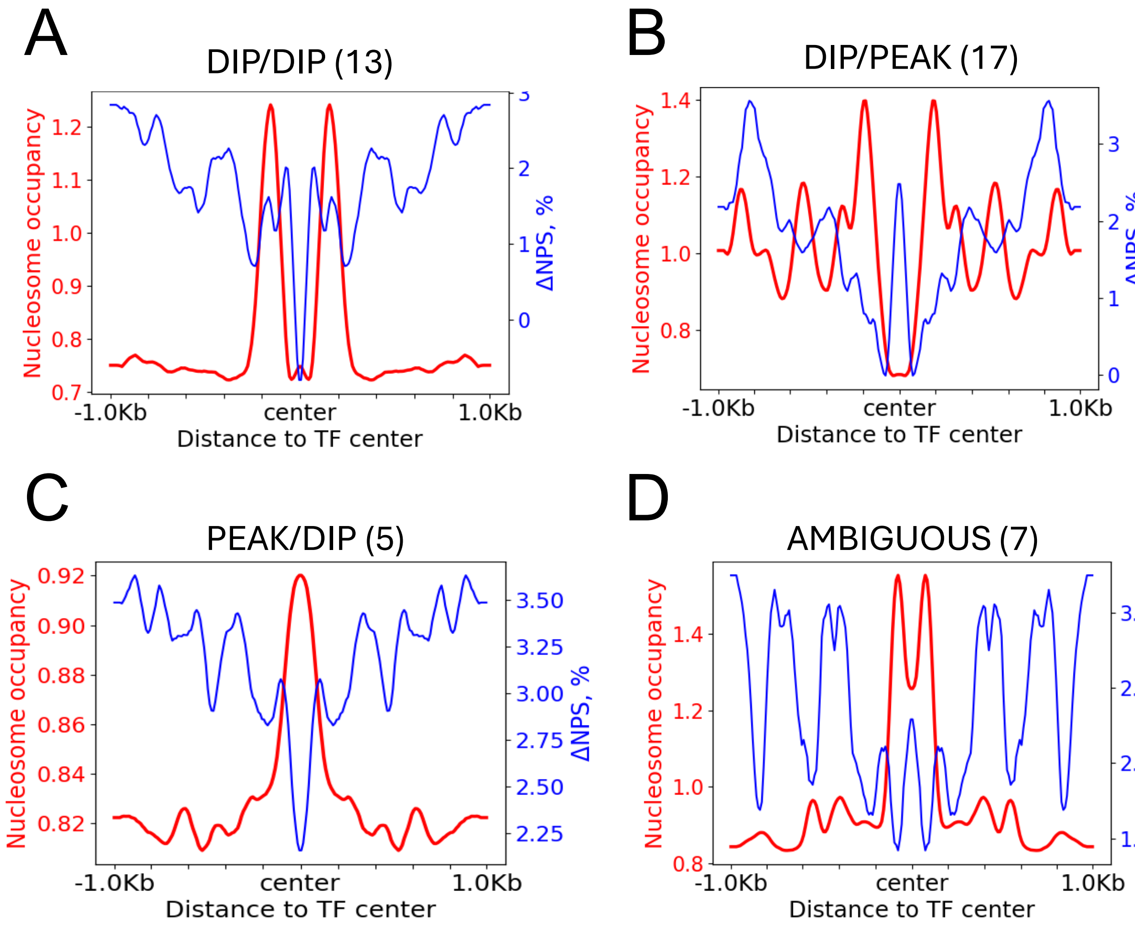

**Figure S10. Average nucleosome occupancy and ΔNPS profiles for TF pattern groups in mouse embryonic stem cells (mESCs).** Average *in vivo* nucleosome occupancy (red) and ΔNPS (blue) profiles are shown for transcription factors in the DIP/DIP (A), DIP/PEAK (B), PEAK/DIP (C), and AMBIGUOUS (D) groups in mESCs. No TFs in this cell line exhibit the PEAK/PEAK pattern. The number of TFs in each group is indicated. Plot notation follows that of Figure 5.

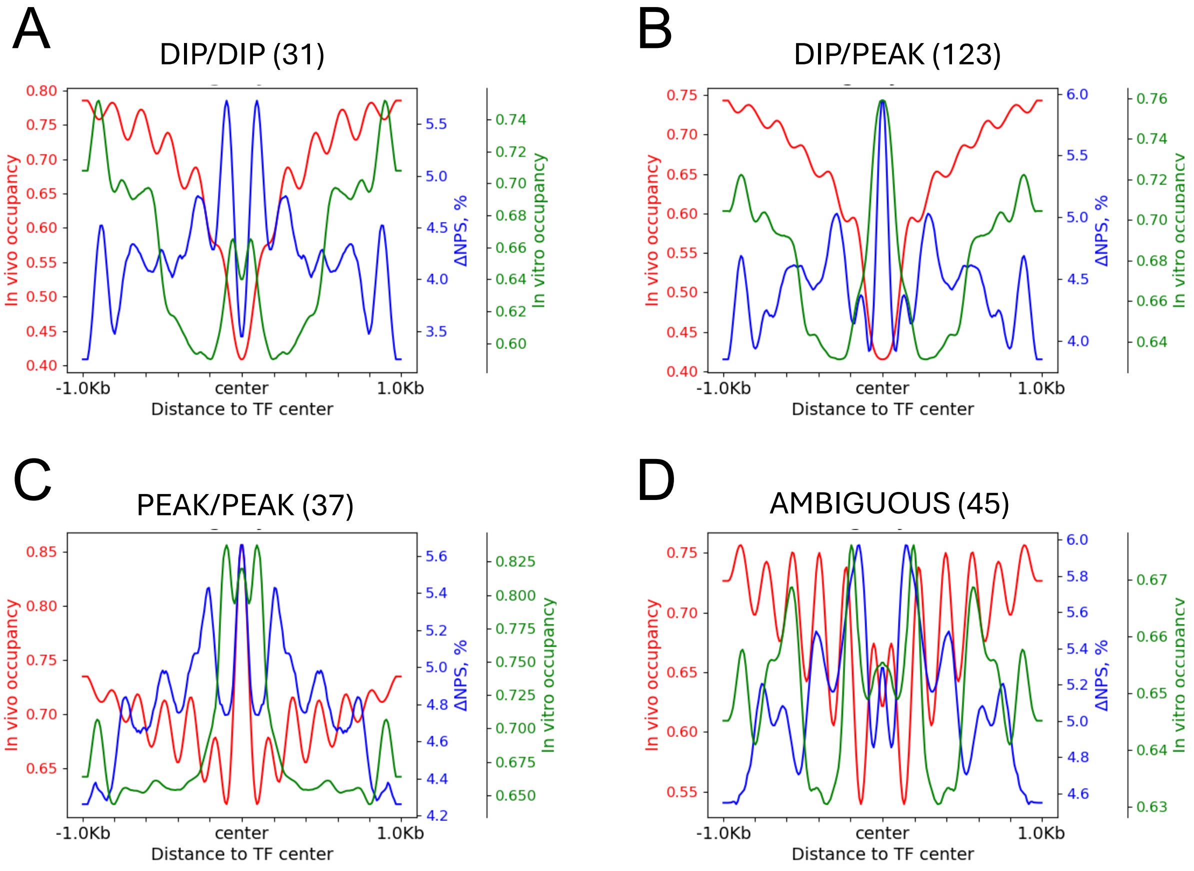

**Figure S11. Average nucleosome occupancy and ΔNPS profiles for TF pattern groups in yeast.** Average *in vivo* nucleosome occupancy (red), *in vitro* nucleosome occupancy (green), and ΔNPS (blue) profiles are shown for transcription factors in the DIP/DIP (A), DIP/PEAK (B), PEAK/PEAK (C), and AMBIGUOUS (D) groups in yeast. No TFs in this organism exhibit the PEAK/DIP pattern. The number of TFs in each group is indicated. Plot notation follows that of Figure 5.

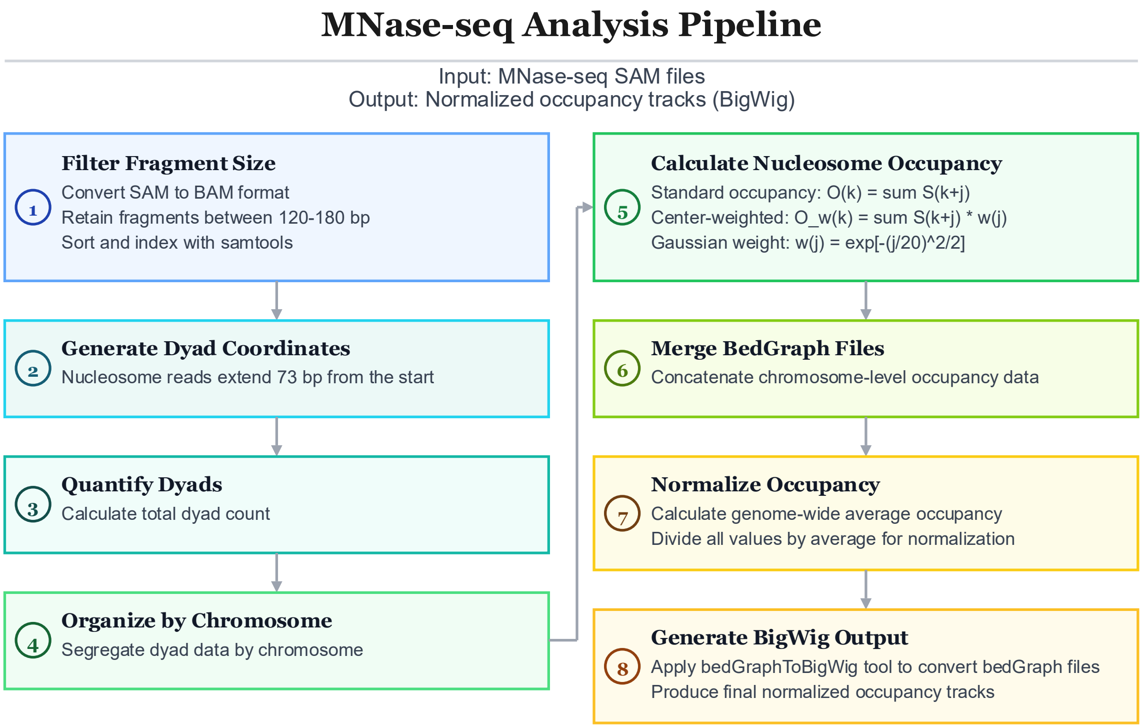

**Figure S12. Nucleosome Occupancy Pipeline:** Data Preprocessing (Steps 1-2) encompasses fragment filtering and dyad coordinate generation from aligned reads. Quantification (Steps 3-4) involves dyad counting and organization by chromosome. Occupancy Analysis (Steps 5-6) calculates nucleosome occupancy using standard and center-weighted methods, followed by merging of chromosome-level bedGraph files. Output Generation (Steps 7-8) performs genome-wide normalization and produces final BigWig tracks for downstream analysis.

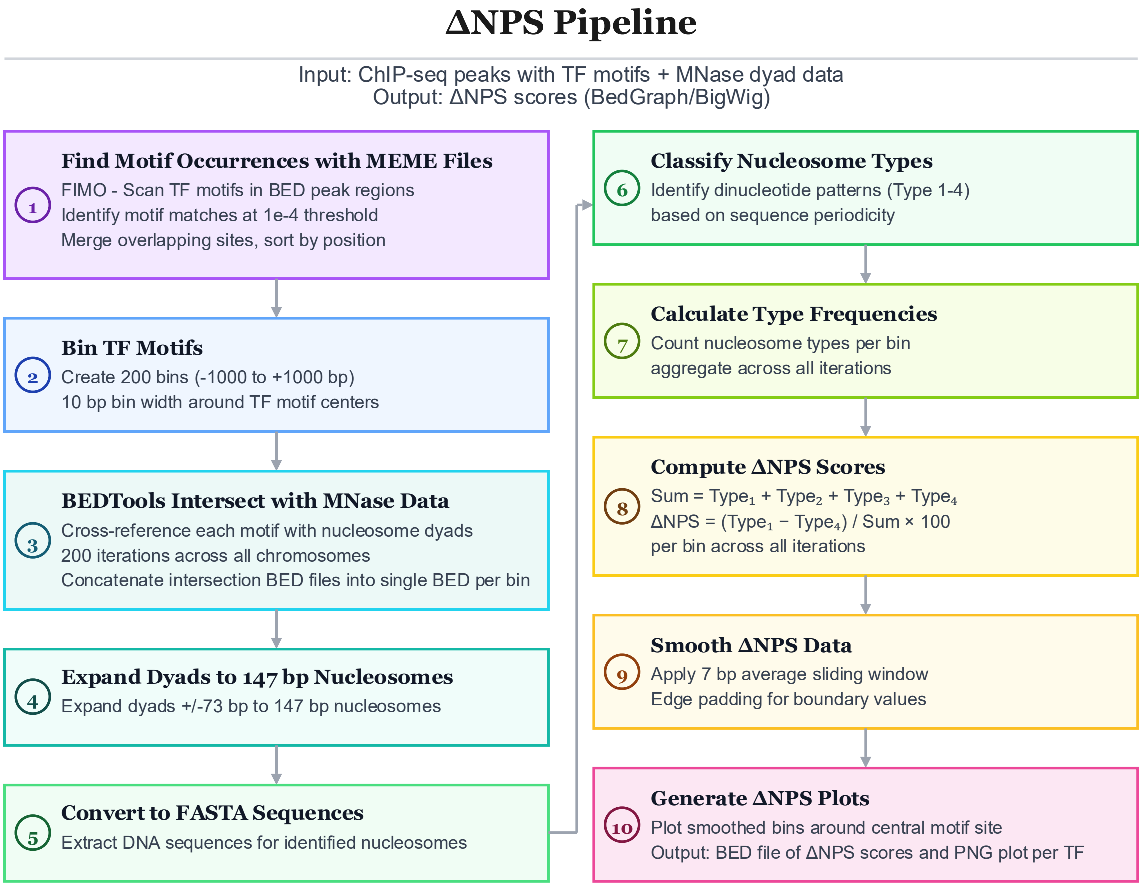

**Figure S13. ΔNPS Pipeline:** Motif Identification (Steps 1-2) discovers transcription factor motif occurrences and organizes them into spatial bins around motif centers. Data Integration (Steps 3-5) intersects motif bins with MNase-seq data, expands dyad positions to average nucleosome size, and extracts underlying DNA sequences. Classification and Quantification (Steps 6-7) assign nucleosomes to rotational positioning types based on dinucleotide periodicity and calculates type frequencies per bin. Score Computation (Steps 8-9) derives ΔNPS values from type distributions and applies smoothing to reduce noise. Visualization (Step 10) generates final plots and output files for publication.
